# Altered Subgenomic RNA Expression in SARS-CoV-2 B.1.1.7 Infections

**DOI:** 10.1101/2021.03.02.433156

**Authors:** Matthew D Parker, Benjamin B. Lindsey, Dhruv R Shah, Sharon Hsu, Alexander J Keeley, David G Partridge, Shay Leary, Alison Cope, Amy State, Katie Johnson, Nasar Ali, Rasha Raghei, Joe Heffer, Nikki Smith, Peijun Zhang, Marta Gallis, Stavroula F Louka, Hailey R Hornsby, Max Whiteley, Benjamin H Foulkes, Stella Christou, Paige Wolverson, Manoj Pohare, Samantha E Hansford, Luke R Green, Cariad Evans, Mohammad Raza, Dennis Wang, Silvana Gaudieri, Simon Mallal, The COVID-19 Genomics UK (COG-UK) consortium, Thushan I. de Silva

## Abstract

SARS-CoV-2 lineage B.1.1.7 viruses are more transmissible, may lead to greater clinical severity, and result in modest reductions in antibody neutralization. subgenomic RNA (sgRNA) is produced by discontinuous transcription of the SARS-CoV-2 genome and is a crucial step in the SARS-CoV-2 life cycle. Applying our tool (periscope) to ARTIC Network Oxford Nanopore genomic sequencing data from 4400 SARS-CoV-2 positive clinical samples, we show that normalised sgRNA expression profiles are significantly increased in B.1.1.7 infections (n=879). This increase is seen over the previous dominant circulating lineage in the UK, B.1.177 (n=943), which is independent of genomic reads, E gene cycle threshold and days since symptom onset at sampling. A noncanonical sgRNA which could represent ORF9b is found in 98.4% of B.1.1.7 SARS-CoV-2 infections compared with only 13.8% of other lineages, with a 16-fold increase in median expression. We hypothesise that this is a direct consequence of a triple nucleotide mutation in nucleocapsid (28280:GAT>CAT, D3L) creating a transcription regulatory-like sequence complementary to a region 3’ of the genomic leader. These findings provide a unique insight into the biology of B.1.1.7 and support monitoring of sgRNA profiles in sequence data to evaluate emerging potential variants of concern.

**One Sentence Summary:** The recently emerged and more transmissible SARS-CoV-2 lineage B.1.1.7 shows greater subgenomic RNA expression in clinical infections and enhanced expression of a noncanonical subgenomic RNA near ORF9b.

## Main Text

The recently emerged SARS-CoV-2 lineages B.1.1.7 (20I/501Y.V1 or VOC-202012/01), B.1.351 (20H/501Y.V2 or VOC-202012/02) and P.1 (20J/501Y.V3 or VOC-202101/02) (*1*) have been classified as variants of concern by public health agencies. An increasing body of evidence suggests B.1.1.7 is more transmissible (*2, 3*) and rapidly became the dominant circulating virus in the United Kingdom (UK) during October 2020 to February 2021 (Figure 1A). To date, B.1.1.7 has been reported in 93 countries (https://cov-lineages.org/, 22^nd^ February 2021), with increasing prevalence. Recent data suggest a similar pattern for P.1 in Manaus (*4*), Brazil (*5*), and now 19 other countries (https://cov-lineages.org/, 22nd February 2021), where it appears to be more transmissible than it’s ancestral lineage B.1.1.28 (*4*). Along with B.1.351, these variants of concern harbour functionally important mutations in the SARS-CoV-2 Spike protein, some of which demonstrate evidence of convergent evolution across all three lineages (Supplementary Table S1). Preliminary data (*6, 7*) shows that B.1.1.7 positive diagnostic respiratory samples may have lower cycle threshold (Ct) values, therefore higher viral loads, compared to other lineages. These findings suggest a potential reason for enhanced transmissibility, though they did not account for potential confounders such as days since symptom onset at sampling. Many of these studies also use S gene target failure (SGTF) as a surrogate for the presence of B.1.1.7 (*6*), which might misclassify samples, depending on the prevalence of B.1.1.7 (*8*). Several analyses from community tested cases also suggest increased mortality associated with B.1.1.7 (*8*). Reasons for the potential viral load increase and enhanced mortality are currently unclear (*9*).

**Figure 1.**
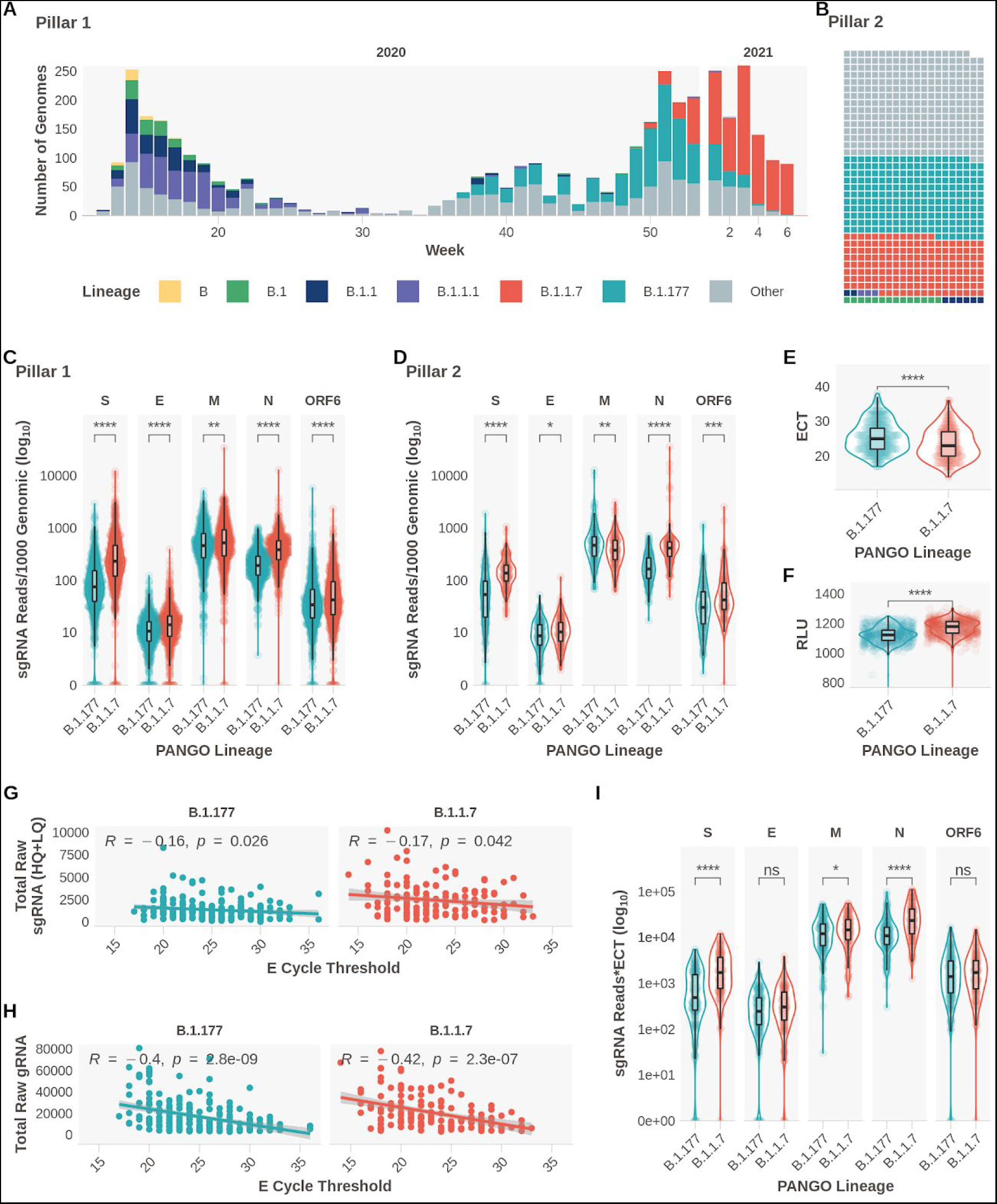
Subgenomic RNA Expression is Increased in B.1.1.7 infections. **A**. Pillar 1 SARS-CoV-2 lineages over time (B.1.1.7_n_ = 729, B.1.177_n_ =764). **B**. Representation of lineage composition of pillar 2 data (B.1.1.7_n_ = 150, B.1.177_n_ = 179). **C**. sgRNA expression in samples of lineages B.1.177 and B.1.1.7 in pillar 1 samples from the most highly expressed ORFs. **D**. sgRNA expression in samples of lineages B.1.177 and B.1.1.7 in pillar 2 samples from the most highly expressed ORFs. **E**. E gene cycle threshold (ECT) for B.1.177 (n=257) and B.1.1.7 (n=185) lineages. **F**. Relative light units (RLU) for B.1.177 (n=626) and B.1.1.7 (n=626) lineages. **G**. E gene cycle threshold compared to total raw sgRNA counts (High and Low Quality) for B.1.177 and B.1.1.7. Correlation coefficient and p value using Pearson. **H**. E gene cycle threshold compared to total raw genomic RNA counts for B.1.177 and B.1.1.7. Correlation coefficient and p value using Pearson. **I**. Raw sgRNA counts normalised to E gene cycle threshold (sgRNA*ECT). All p-values (except G & H) calculated using an unpaired Wilcoxon signed rank test, and adjusted for multiple testing with the Holm method (**** < 0.0001, *** < 0.001, ** < 0.01, * < 0.05)

Genomic surveillance has been critical in rapidly identifying these variants and Nanopore sequencing of ARTIC Network (*10*) prepared SARS-CoV-2 amplicons is used by many laboratories to generate this data. We have recently reported an approach to quantify subgenomic RNA (sgRNA) expression profiles from genomic sequence data, which is produced as a result of a critical step in the SARS-CoV-2 replication cycle (*11*). sgRNA is produced from the genomically encoded SARS-CoV-2 RNA Dependent RNA polymerase (RdRp) using discontinuous transcription of the positive, single-stranded SARS-CoV-2 genome from the 3’ end. Negatively stranded RNAs are produced, which are shorter than the genome, owing to a template switch from the ORF to the leader sequence at the 5’ end of the genome when RdRp encounters a transcription regulatory sequence in the genome body (TRS-B) to a complementary TRS 3’ of the leader sequence (TRS-L). All sgRNAs therefore contain a leader sequence at their 3’ end which can be used computationally for their identification. There are thought to be nine such canonical sgRNAs; Spike:S, E: Envelope, M: Membrane, N: Nucleocapsid, ORF3a, ORF6, ORF7a, ORF8 and ORF10, although multiple studies have found negligible ORF10 expression (*12, 13*).

As part of COVID-19 Genomics Consortium UK (COG-UK) (*14*) we are sequencing SARS-CoV-2 positive nose-throat swabs from healthcare workers and patients at Sheffield Teaching Hospitals NHS Foundation Trust, Sheffield, UK (‘Pillar 1’ testing). Additionally, to relieve pressure on centralised sequencing services, we also sequence a selection from ‘Pillar 2’ testing, which represent SARS-CoV-2 positive samples from the community, tested at the UK’s Lighthouse Laboratories (Figure S1). We hypothesised that we would see differences in sgRNA expression profiles in distinct lineages of SARS-CoV-2, in particular, increased sgRNA in B.1.1.7 that may relate to its altered phenotype.

We stratified sgRNA expression by lineage in 4400 SARS-CoV-2 sequences that reached our previously defined quality control thresholds (>90% genome coverage, >50K mapped reads, Table S2), normalized for the genomic coverage from the corresponding amplicon for the five most abundantly expressed open reading frames (S, E, M, N and ORF6). We focused our analyses primarily on comparing B.1.1.7 with B.1.177, the dominant lineage in the UK, which was displaced by B.1.1.7. These two lineages are also the most represented sequences from both sampling pillars in our dataset (Figure 1A&B, B.1.1.7; Pillar 1: 729, Pillar 2: 150, B.1.177; Pillar 1: 764 Pillar 2: 179). A significant increase in normalized sgRNA expression of S, N, and to a lesser extent E, ORF6 and M, was apparent in B.1.1.7 SARS-CoV-2 infections (Figure 1C&D, Wilcoxon effect sizes; S: 0.475, E: 0.191, M 0.0700, N: 0.469, ORF6: 0.105). Negligible differences were seen in other ORF sgRNA (Figure S2). Consistent with previous findings (*6, 15*), we also found significantly decreased E gene Ct (ECT; in house diagnostic assay (*16*); B.1.177 median=25, B.1.1.7 median=23) and significantly greater Relative Light Units (RLU; Hologic Panther platform; B.1.177 median=1121, B.1.1.7 median = 1177) in B.1.1.7 infections compared to B.1.177 infections (Figure 1E&F). Accordingly, total mapped reads and total genomic RNA reads (i.e. non-sgRNA) were also significantly higher for B.1.1.7 sequences compared to B.1.177 (Figure S3).

We observed a weak but significant negative correlation between raw sgRNA reads and ECT (R = -0.16 and p=0.026 for B.1.177, R = -0.17 and p=0.042 for B.1.1.7), with a greater negative correlation between genomic reads and ECT (R = -0.4 and p<0.0001 for B.1.177, R = -0.42 and p<0.0001 for B.1.1.7; Fig 1G&H). To ensure the observed sgRNA expression increase in B.1.1.7 was not due to the effect of ECT alone (as a surrogate of viral load), we normalized sgRNA to ECT and repeated our comparisons. The differences between B.1.1.7 and B.1.177 were still apparent for S, N and M sgRNA (Figure 1I), suggesting that the increase in B.1.1.7 sgRNA expression is seen independently of any difference in ECT between lineages. To ensure the changes observed were not due to parental lineage changes in sgRNA expression such as those observed with the R203K/G204R mutation (*17*) we analysed normalized sgRNA comparing B.1.1.7 and B.1.177 with their respective ancestral lineages (Figure 2A&B). An increase in sgRNA is still observed for B.1.1.7 compared to B.1.1 in all major ORFs and interestingly, for B.1.177 compared to B.1 for ORFs S, E, M and N (Figure 2A&B). This demonstrates the importance of stratifying sequences by lineage when studying sgRNA.

**Figure 2.**
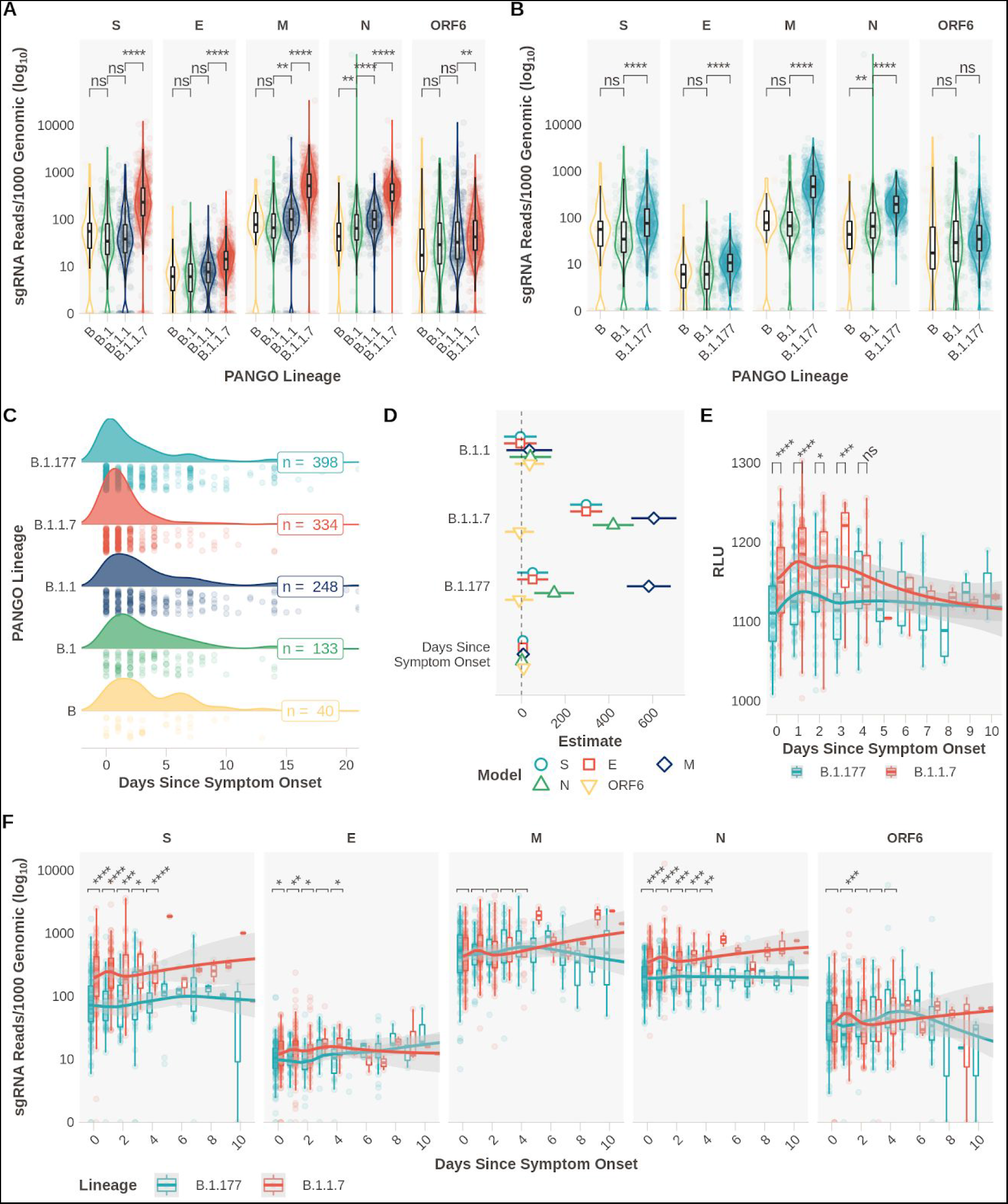
Relationship between day of symptom onset at sampling and subgenomic RNA levels in B.1.1.7 and B.1.177 infections. **A**. Subgenomic RNA data from most highly expressed ORFs for B.1.1.7 and respective ancestral lineages in Pillar 1 data (B=_n_51, B.1=_n_164, B.1.1_n_=302,,B.1.1.7_n_=729). **B**. Subgenomic RNA data from most highly expressed ORFs for B.1.177 and respective ancestral lineages in Pillar 1 data (B_n_=51, B.1=164, B.1.177_n_=764). **C**. Distribution of the reported days since symptom at the time of sampling for B (n=40), B.1 (n=133), B.1.1 (n=248), B.1.177 (n=398) and B.1.1.7 (n=334) lineage infections. **D**. Model estimates and 95% confidence intervals from generalized linear model comparing effect of lineage and day of symptom onset on sgRNA expression for mostly highly abundant ORFs. Reference lineage B.1. No interaction was observed between any lineage and day of symptom onset. One B.1 sample with outlier N expression (SHEF-CC370) was excluded. **E**. Pseudo time course of RLU values for B.1.1.7 and B.1.177, plot restricted to 10 days. (0 Days; B.1.177_n_ = 92, B.1.1.7_n_= 106. 1 Day; B.1.177_n_ = 52, B.1.1.7_n_= 102. 2 Days; B.1.177_n_ = 28, B.1.1.7_n_= 43. 3 Days; B.1.177_n_ = 31, B.1.1.7_n_= 14. 4 Days; B.1.177_n_ = 17, B.1.1.7_n_= 19.). **F**. Normalized subgenomic RNA expression for each of the highest expressed ORFs stratified by the reported days since symptom onset, loess model. Note: differing y-axis scales, plot restricted to 10 days. (0 Days; B.1.177_n_ = 115, B.1.1.7_n_= 93. 1 Day; B.1.177_n_ = 53, B.1.1.7_n_= 93. 2 Days; B.1.177_n_ = 31, B.1.1.7_n_= 47. 3 Days; B.1.177_n_ = 31, B.1.1.7_n_= 12. 4 Days; B.1.177_n_ = 19, B.1.1.7_n_= 18.). All p-values calculated using an unpaired Wilcoxon signed rank test, and adjusted for multiple testing with Holm (**** < 0.0001, *** < 0.001, ** < 0.01, * < 0.05, ns or blank = not significant).

It is possible that the days since symptom onset at sampling may vary between lineages in our dataset, either due to changes in sampling practice over time or presentation of individuals to healthcare services, which in turn could confound our sgRNA findings. We obtained information on symptom duration at sampling for 2327 samples in our pillar 1 genome sequenced cohort. A modest difference in the distribution of the days since symptom onset between B.1.177 and B.1.1.7 was apparent, where B.1.1.7 infections appeared to be sampled earlier (Figure 2C). We therefore applied a generalized linear regression model to adjust for any effect this change in distribution might have on sgRNA expression levels in lineages B.1, B.1.1, B.1.177 and B.1.1.7. Subgenomic RNA was still significantly elevated in B.1.1.7 in S, N and M and to a lesser extent in E (Figure 2D, Fig S4, Tables S3&S4). A smaller increase in sgRNA in B.1.177 (compared to B.1) was still seen, though only for M and N. Furthermore, as sampling practice and sequencing in healthcare workers in our hospital remained consistent throughout the period when B.1.177 and B.1.1.7 were circulating, we hypothesized that these data were less likely to be affected by unknown confounders in sampling. Subgenomic RNA levels remained significantly higher in B.1.1.7 compared to B.1.177 infections when the comparison was restricted to samples from healthcare workers (Figure S5). No difference in sgRNA levels for B.1.1.7 was seen by sex or age (Figure S6 and S7). Relative light units in the diagnostic assay also varied over time in B.1.1.7 compared to B.1.177 infections, with significantly higher RLU at 0, 1, 2, 3 days since symptom onset (Figure 2E). Data since symptom onset for samples with Ct values were insufficient to replicate this analysis.

From these cross-sectional data we created a pseudo infection time course using normalized sgRNA expression at each day following symptom onset between B.1.1.7 and B.1.177 infections. Significantly higher sgRNA for B.1.1.7 compared to B.1.177 was observed early in infection for S and N sgRNA, with lower magnitude differences for E sgRNA (Figure 2F&S8). This increase appeared to be greatest on day one following symptom onset. Using the same methods for sgRNA estimation, we have previously shown that *ACE2* and *TMPRSS2* expression alter the kinetics of sgRNA expression *in vitro*, leading to a peak in expression at an earlier stage of infection (*11*). It is possible that the increased affinity for *ACE2* conferred by N501Y in the B.1.1.7 spike protein (*18, 19*) enhances sgRNA profiles early in clinical infection. Experimental validation of these findings in cell and animal models would be extremely valuable.

Noncanonical sgRNA are the result of RdRp template switching from regions of the genome with no canonical TRS-B site. These noncanonical sgRNA tend to be enriched around canonical sites, presumably due to the frequency of RdRp template switching that occurs in close proximity. We quantified the expression of these noncanonical sgRNA in pillar 1 B.1.1.7 infections. We observed low levels of these transcripts throughout the genome across all samples (*11*) and compared the presence of noncanonical sgRNA transcripts between B.1.1.7 and B.1.177 infections (Figure 3A). In addition to the noncanonical sgRNA resulting from the nucleocapsid R203K/G204R mutation (N*, Figure S2) (*17*), B.1.1.7 samples were also significantly enriched for reads which support the production of a noncanonical sgRNA from genomic position 28282 (Figure 3B,C&D, Fig S9, Wilcoxon unpaired p = <2e-16, Wilcoxon effect size = 0.797, B.1.1.7_n_=717, other_n_=430). The B.1.1.7 genomic sequence contains a triple nucleotide mutation at 28280, 28281, and 28282 GAT>CTA, resulting in an amino acid substitution, D3L, in the nucleocapsid protein. This triplet results in enhanced complementarity between the 28282 genomic region and the sequence 3’ of the leader at the 5’ end of the genome (Figure 3E), which may have resulted in a novel TRS-B-like site that drives higher sgRNA production from this locus. Of note, this site is upstream of the ORF9b ATG and this noncanonical sgRNA retains the full coding region of the ORF9b, but lacks the canonical start codon of N. We propose that this represents the ORF9b sgRNA, which was detected at low levels in 13.8% (430/3127) non-B.1.1.7 samples (median normalised expression=0.32), but is found in almost all pillar 1 B.1.1.7 infections (98.4%, 717/729), with a 16-fold increase in expression (median normalised expression=6.02, Figure 3A&C).

**Figure 3.**
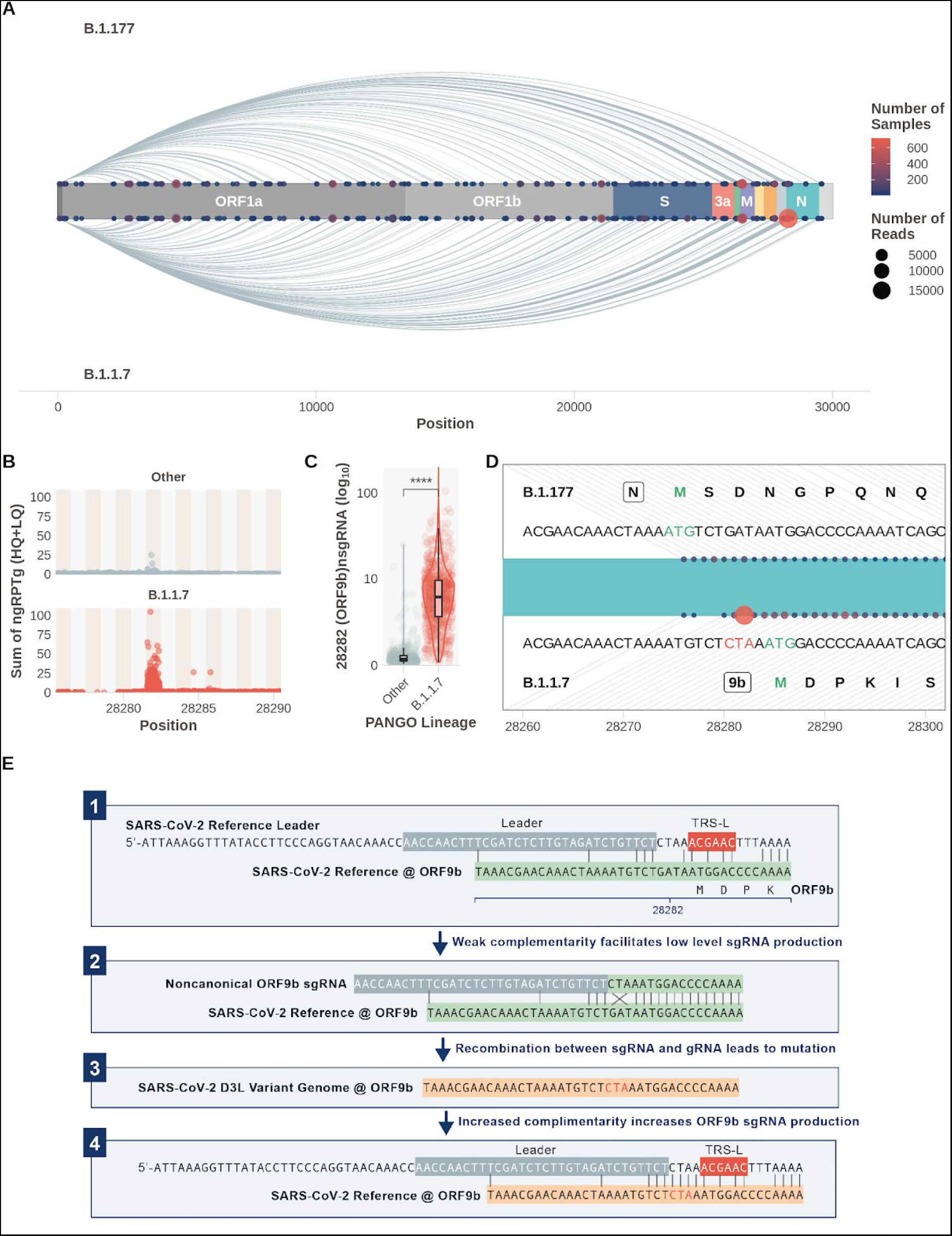
A Noncanonical sgRNA representing ORF9b is highly expressed in B.1.1.7 due to a triple nucleotide mutation in nucleocapsid leading to the D3L substitution. **A**. Subgenomic RNA not attributed to a canonical TRS-B site in B.1.177 (top) and B.1.1.7 (bottom) SARS-CoV-2 infections (Pillar 1 data). Size of point is number of reads, colour is number of samples (N* excluded for clarity). **B**. Total normalized noncanonical sgRNA (ngRPTg) in B.1.1.7 vs other lineages. **C**. B.1.1.7 has significantly increased expression of a noncanonical sgRNA at 28282. (Wilcoxon effect size, unpaired = 0.797, Other_n_=430, B.1.1.7_n_=717). **D**. Schematic of the noncanonical sgRNA (points) in the context of the SARS-CoV-2 genome around position 28282. Top shows the sequence present in B.1.177 and the amino acid sequence of the N protein. Bottom shows the sequence present in B.1.1.7 with the triplet CTA mutation and the closest ATG which represents the ORF9b methionine. **E**. Proposed model for sgRNA driven mutation of N D3L which leads to high ORF9b expression in B.1.1.7. All p-values calculated using an unpaired Wilcoxon signed rank test, and adjusted for multiple testing with holm (**** < 0.0001, *** < 0.001, ** < 0.01, * < 0.05, ns = not significant)

Interestingly, we noticed non-B.1.1.7 samples in our dataset with low levels of putative ORF9b sgRNA did not contain the nucleocapsid D3L mutation. We propose the following hypothesis to explain our findings: in non-B.1.1.7 SARS-CoV-2, the weak complementarity around the ORF9b ATG causes low levels of noncanonical sgRNA to be produced which has high complementarity to the genome, but crucially contains the CTA triplet (Figure S9A). This could have led to a transcriptionally driven recombination event resulting in the GAT > CTA mutation in the genomic sequence (Figure 3E panel 2&3). This new TRS-B-like site with increased complementarity in B.1.1.7 viruses, drives higher sgRNA production from this locus, akin to sites of canonical sgRNA production (Figure 3E Panel 4). A similar event has been described previously more 3’ in the N ORF (*17*), and mutations which result in increased complementarity and noncanonical sgRNA formation have been shown to occur in coronaviruses (*20*). This raises the possibility that an ORF9b protein could be produced independently of other upstream sgRNAs with greater efficiency in B.1.1.7. ORF9b has been shown to regulate interferon responses (*21*–*23*) and has been shown to be present in SARS-CoV (*24*) and SARS-CoV-2 infections (*13, 24, 25*). This finding could therefore be important for investigating the greater transmissibility of B.1.1.7 and requires urgent experimental validation. We explored whether the nucleocapsid D3L mutation has occurred in non-B.1.1.7 lineages in the global COG-UK sequence data set (14th of February 2021). 77 sequences with D3L were noted in a variety of SARS-CoV-2 lineages and appear as homoplasies in a global phylogeny, suggesting that this event may have occurred independently on several occasions (Figure S10).

Taken together, our data suggests that sgRNA expression from existing ARTIC Nanopore sequencing data can be used in real time to examine the effect of SARS-CoV-2 variation on its ability to express its genome. We cannot say if the increased sgRNA expression is the cause or consequence of an increase in viral replication or a more efficacious entry, but our study provides further insight to guide exploration with mechanistic studies. A major advantage of this approach is that we can deconvolute the contribution of genomic and subgenomic RNA, which is impossible with current diagnostic PCR assays and we can, additionally, examine all ORFs simultaneously and discover noncanonical subgenomic RNA which could be of biological relevance. Finally we believe that sgRNA expression analysis should be carried out on all compatible genomic surveillance platforms to give an instant readout of altered expression profiles in emerging SARS-CoV-2 variants. This would use existing data to complement epidemiological and phylodynamic methods, and provide an early warning of variants that might be of concern with regards to greater transmissibility and/or disease severity.

## Methods

### Diagnostic SARS-CoV-2 Testing

SARS-CoV-2 positivity was determined from nose/throat swabs diagnostically by Sheffield Teaching Hospitals NHS Foundation trust either using the Hologic Panther to generate Relative Light Units (RLU) (*26*) or an in house dual E/RdRp real time PCR assay to generate a cycle threshold (ECT or RCT respectively) (*16*).

### Sample Preparation, ARTIC Network PCR and Nanopore Sequencing

RNA was extracted from viral transport medium (VTM) with Qiagen QIAamp MinElute Virus Spin Kit (50). Resultant RNA was subject to the ARTIC network (*10*) tiled amplicon protocol and subsequently sequenced on an Oxford Nanopore GridION X5. Bases were called with the default basecaller in MinKNOW (currently guppy v4) with *--require-both-ends* set for de-multiplexing. Raw sequencing fastq files were subject to no processing after sequencing to ensure sgRNA leader sequences are retained as described previously (*11*).

### Subgenomic Read Classification and Normalisation

Subgenomic RNAs were classified using periscope (*11*) as described previously. Briefly, reads containing the leader sequence at their start are identified by a local alignment, the quality of this alignment determines which quality “bin” sgRNA reads are placed (HQ, LQ or LLQ). The amplicon from which the sgRNA evidence was generated is determined and a count of genomic reads for this amplicon used to normalise the raw sgRNA read counts.

Samples were excluded from the subsequent analysis if:

- Their consensus coverage was < 0.9
- They had less than 50,000 mapped reads (we have previously shown that fewer reads produce a less robust analysis)

### Lineage Assignment

Lineages (*1*) were assigned using Pangolin (https://github.com/cov-lineages/pangolinv2.1.7).

### Figure Generation, Statistical Analysis

Statistical analysis was performed in R (*27*). Figures were generated in R using tidyverse (*28*)apart from those that depict sequencing reads, which were generated in IGV (*29*). Tests between groups were performed using an unpaired Wilcoxon test using the rstatix package (https://github.com/kassambara/rstatix), adjusting p-values for any multiple comparisons using the Holm method.

### Phylogenetic Tree Generation

The grapevine pipeline (https://github.com/COG-UK/grapevine) was used for generating the phylogeny based on all data available on GISAID and COG-UK up until 16th February 2021. A representative sample of global sequences was obtained in 2 steps. First by randomly selecting one sequence per country per epi week followed by random sampling of the remaining sequences to generate a sample of 4000 sequences. The global tree was then pruned using code adapted from the tree-manip package (https://github.com/josephhughes/tree-manip). We then identified samples with D3L mutations and colour coded these tips according to their lineages. The visualisation was produced using R/ape, R/ggplot2, R/ggtree, R/treeio, R/phangorn, R/stringr, R/dplyr, R/aplot.

## Ethics Approval and Consent

Individuals presenting with active COVID-19 disease were sampled for SARS CoV-2 sequencing at Sheffield Teaching Hospitals NHS Foundation Trust, UK using samples collected for routine clinical diagnostic use. This work was performed under approval by the Public Health England Research Ethics and Governance Group for the COVID-19 Genomics UK consortium (R&D NR0195).

## Data Availability

All SARS-CoV-2 consensus sequences that are of high enough quality are available on GISAID and ENA and from https://www.cogconsortium.uk/data/. All sgRNA expression data is provided as supplementary files and at https://github.com/sheffield-bioinformatics-core/periscope-variants-publication. All raw sequencing data will be available on ENA as soon as possible.

## Code Availability

Periscope is available at https://github.com/sheffield-bioinformatics-core/periscope and the code used to generate the figures contained within this manuscript can be found as a supplementary files or at https://github.com/sheffield-bioinformatics-core/periscope-variants-publication

## Acknowledgements

We thank the Sheffield Bioinformatics Core for their useful thoughts and discussions. We would like to thank the members of the Sheffield Biomedical Research centre for their continued support of the SARS-CoV-2 sequencing work in Sheffield.

## Author Contributions

**Contributor Role* Contributor**

Conceptualization MDP, TIdS

Data curation MDP, TIdS, BL, DS, AK, DP, SH, MR, CE

Formal Analysis MDP, TIdS, BL, SH

Funding acquisition DW, TIdS

Investigation PZ, MG, SL, MW, BF, SC, PW, MP, LG

Methodology MDP, TIdS

Project administration NS, SH

Resources CE, MR, DP, AC, AS, KJ, NA, RR

Software MDP, JH

Supervision MDP, DW, TIdS

Validation Not Applicable

Visualization MDP, BL, SH

Writing – original draft MDP, TIdS

Writing – review & editing MDP, DW, BL, SH, TIdS

*Contributions as described at: https://casrai.org/credit/

## Funding

Sequencing of SARS-CoV-2 samples was undertaken by the Sheffield COVID-19 Genomics Group as part of the COG-UK CONSORTIUM and supported by funding from the Medical Research Council (MRC) part of UK Research & Innovation (UKRI), the National Institute of Health Research (NIHR) and Genome Research Limited, operating as the Wellcome Sanger Institute. MDP and DW are funded by the NIHR Sheffield Biomedical Research Centre (BRC - IS-BRC-1215-20017). TIdS is supported by a Wellcome Trust Intermediate Clinical Fellowship (110058/Z/15/Z). We thank all partners of and contributors to the COG-UK consortium, who are listed at https://www.cogconsortium.uk/about/.

## Competing Interests

The authors note no competing interests.

## Notes

### Competing Interest Statement

The authors have declared no competing interest.

### Summary of Updates

Update to the abstract to match the PDF.

