## Supplementary Material for "Altered Subgenomic RNA Expression in SARS-CoV-2 B.1.1.7 Infections"

#### Contents

**Supplementary Material** **1**
Supplementary Authors 2
Supplementary Files 10
Supplementary Tables 11
Supplementary Figures 15

### Supplementary Authors

**Funding acquisition, Leadership and supervision, Metadata curation, Project administration, Samples and logistics, Sequencing and analysis, Software and analysis tools, and Visualisation:**
Dr Samuel C Robson PhD [13].

**Funding acquisition, Leadership and supervision, Metadata curation, Project administration, Samples and logistics, Sequencing and analysis, and Software and analysis tools:**
Prof Nicholas J Loman PhD [41] and Dr Thomas R Connor PhD [10, 69].

**Leadership and supervision, Metadata curation, Project administration, Samples and logistics, Sequencing and analysis, Software and analysis tools, and Visualisation:**
Dr Tanya Golubchik PhD [5].

**Funding acquisition, Metadata curation, Samples and logistics, Sequencing and analysis, Software and analysis tools, and Visualisation:**
Dr Rocio T Martinez Nunez PhD [42].

**Funding acquisition, Leadership and supervision, Metadata curation, Project administration, and Samples and logistics:**
Dr Catherine Ludden PhD [88].

**Funding acquisition, Leadership and supervision, Metadata curation, Samples and logistics, and Sequencing and analysis:**
Dr Sally Corden PhD [69].

**Funding acquisition, Leadership and supervision, Project administration, Samples and logistics, and Sequencing and analysis:**
Ian Johnston [99] and Dr David Bonsall PhD [5].

**Funding acquisition, Leadership and supervision, Sequencing and analysis, Software and analysis tools, and Visualisation:**
Prof Colin P Smith PhD [87] and Dr Ali R Awan PhD [28].

**Funding acquisition, Samples and logistics, Sequencing and analysis, Software and analysis tools, and Visualisation:**
Dr Giselda Bucca PhD [87].

**Leadership and supervision, Metadata curation, Project administration, Samples and logistics, and Sequencing and analysis:**
Dr M. Estee Torok FRCP [22, 101].

**Leadership and supervision, Metadata curation, Project administration, Samples and logistics, and Visualisation:**
Dr Kordo Saeed MD/ FRCPath [81, 110] and Dr Jacqui A Prieto PhD [83, 109].

**Leadership and supervision, Metadata curation, Project administration, Sequencing and analysis, and Software and analysis tools:**
Dr David K Jackson PhD [99].

**Metadata curation, Project administration, Samples and logistics, Sequencing and analysis, and Software and analysis tools:**
Dr William L Hamilton PhD [22].

**Metadata curation, Project administration, Samples and logistics, Sequencing and analysis, and Visualisation:**
Dr Luke B Snell MSc/ MBBS [11].

**Funding acquisition, Leadership and supervision, Metadata curation, and Samples and logistics:**
Dr Catherine Moore [69].

**Funding acquisition, Leadership and supervision, Project administration, and Samples and logistics:**
Dr Ewan M Harrison PhD [99, 88].

**Leadership and supervision, Metadata curation, Project administration, and Samples and logistics:**
Dr Sonia Goncalves PhD [99].

**Leadership and supervision, Metadata curation, Samples and logistics, and Sequencing and analysis:**
Prof Ian G Goodfellow PhD [24], Dr Derek J Fairley PhD [3, 72], Prof Matthew W Loose PhD [18] and Joanne Watkins MSc [69].

**Leadership and supervision, Metadata curation, Samples and logistics, and Software and analysis tools:**
Rich Livett MSc [99].

**Leadership and supervision, Metadata curation, Samples and logistics, and Visualisation:**
Dr Samuel Moses MD [25, 106].

**Leadership and supervision, Metadata curation, Sequencing and analysis, and Software and analysis tools:**
Dr Roberto Amato PhD [99], Dr Sam Nicholls PhD [41] and Dr Matthew Bull PhD [69].

**Leadership and supervision, Project administration, Samples and logistics, and Sequencing and analysis:**
Prof Darren L Smith PhD [37, 58, 105].

**Leadership and supervision, Sequencing and analysis, Software and analysis tools, and Visualisation:**
Dr Jeff Barrett PhD [99] and Prof David M Aanensen PhD [14, 114].

**Metadata curation, Project administration, Samples and logistics, and Sequencing and analysis:**
Dr Martin D Curran PhD [65], Dr Surendra Parmar PhD [65], Dr Dinesh Aggarwal MRCP [95, 99, 64] and Dr James G Shepherd MBChB/MRCP [48].

**Metadata curation, Project administration, Sequencing and analysis, and Software and analysis tools:**
Dr Matthew D Parker PhD [93].

**Metadata curation, Samples and logistics, Sequencing and analysis, and Visualisation:**
Dr Sharon Glaysher PhD [61].

**Metadata curation, Sequencing and analysis, Software and analysis tools, and Visualisation:**

Dr Matthew Bashton PhD [37, 58], Dr Anthony P Underwood PhD [14, 114], Dr Nicole Pacchiarini PhD [69] and Dr Katie F Loveson PhD [77].

**Project administration, Sequencing and analysis, Software and analysis tools, and Visualisation:**
Dr Alessandro M Carabelli PhD [88].

**Funding acquisition, Leadership and supervision, and Metadata curation:**
Dr Kate E Templeton PhD [53, 90].

**Funding acquisition, Leadership and supervision, and Project administration:**
Dr Cordelia F Langford PhD [99], John Sillitoe BEng [99], Dr Thushan I de Silva PhD [93] and Dr Dennis Wang PhD [93].

**Funding acquisition, Leadership and supervision, and Sequencing and analysis:**
Prof Dominic Kwiatkowski [99, 107], Prof Andrew Rambaut DPhil [90], Dr Justin O'Grady PhD [70, 89] and Dr Simon Cottrell PhD [69].

**Leadership and supervision, Metadata curation, and Sequencing and analysis:**
Prof Matthew T.G. Holden PhD [68] and Prof Emma C Thomson PhD/FRCP [48].

**Leadership and supervision, Project administration, and Samples and logistics:**
Dr Husam Osman PhD [64, 36], Dr Monique Andersson PhD [59], Prof Anoop J Chauhan [61] and Dr Mohammed O Hassan-Ibrahim PhD/FRCPath [6].

**Leadership and supervision, Project administration, and Sequencing and analysis:**
Dr Mara Lawniczak [99].

**Leadership and supervision, Samples and logistics, and Sequencing and analysis:**
Prof Ravi Kumar Gupta PhD [88, 113], Dr Alex Alderton PhD [99], Dr Meera Chand [66], Dr Chrystala Constantinidou PhD [94], Dr Meera Unnikrishnan PhD [94], Prof Alistair C Darby PhD [92], Prof Julian A Hiscox PhD [92] and Prof Steve Paterson PhD [92].

**Leadership and supervision, Sequencing and analysis, and Software and analysis tools:**
Dr Inigo Martincorena [99], Prof David L Robertson PhD [48], Dr Erik M Volz PhD [39], Dr Andrew J Page PhD [7] and Prof Oliver G Pybus DPhil [23].

**Leadership and supervision, Sequencing and analysis, and Visualisation:**
Dr Andrew R Bassett PhD [99].

**Metadata curation, Project administration, and Samples and logistics:**
Dr Cristina V Ariani PhD [99], Dr Michael H Spencer Chapman MBBS [99, 88], Dr Kathy K Li MBBCh/FRCPath [48], Dr Rajiv N Shah BMBS/MRCP/MSc [48], Dr Natasha G Jesudason MBChB MRCP FRCPath [48] and Dr Yusri Taha MD/PhD [50].

**Metadata curation, Project administration, and Sequencing and analysis:**
Martin P McHugh MSc [53], Dr Rebecca Dewar PhD [53].

**Metadata curation, Samples and logistics, and Sequencing and analysis:**
Dr Aminu S Jahun PhD [24], Dr Claire McMurray PhD [41], Ms Sarojini Pandey MSc [84], Dr James P McKenna PhD [3], Dr Andrew Nelson PhD [58, 105], Dr Gregory R Young PhD [37, 58], Dr Clare M McCann PhD [58, 105] and Mr Scott Elliott [61].

**Metadata curation, Samples and logistics, and Visualisation:**
Ms Hannah Lowe MSc [25].

**Metadata curation, Sequencing and analysis, and Software and analysis tools:**

Dr Ben Temperton Ph.D. [91], Dr Sunando Roy PhD [82], Dr Anna Price PhD [10], Dr Sara Rey PhD [69] and Mr Matthew Wyles [93].

**Metadata curation, Sequencing and analysis, and Visualisation:**
Stefan Rooke MSc [90] and Dr Sharif Shaaban PhD [68].

**Project administration, Samples and logistics, Sequencing and analysis:**
Dr Mariateresa de Cesare PhD [98].

**Project administration, Samples and logistics, and Software and analysis tools:**
Laura Letchford BSc [99].

**Project administration, Samples and logistics, and Visualisation:**
Miss Siona Silveira MSc [81], Dr Emanuela Pelosi FRCPath [81] and Dr Eleri Wilson-Davies MD/FRCPath [81].

**Samples and logistics, Sequencing and analysis, and Software and analysis tools:**
Dr Myra Hosmillo PhD [24].

**Sequencing and analysis, Software and analysis tools, and Visualisation:**
Áine O'Toole MSc [90], Dr Andrew R Hesketh PhD [87], Mr Richard Stark MSc [94], Dr Louis du Plessis PhD [23], Dr Chris Ruis PhD [88], Dr Helen Adams PhD [4] and Dr Yann Bourgeois PhD [76].

**Funding acquisition, and Leadership and supervision:**
Dr Stephen L Michell PhD [91], Prof Dimitris Grammatopoulos PhD/FRCPath [84, 112], Dr Jonathan Edgeworth PhD/FRCPath [12], Prof Judith Breuer MD [30, 82], Prof John A Todd PhD [98] and Dr Christophe Fraser PhD [5].

**Funding acquisition, and Project administration:**
Dr David Buck PhD [98] and Michaela John BSc [9].

**Leadership and supervision, and Metadata curation:**
Dr Gemma L Kay PhD [70].

**Leadership and supervision, and Project administration:**
Steve Palmer [99], Prof Sharon J Peacock [88, 64] and David Heyburn [69].

**Leadership and supervision, and Samples and logistics:**
Danni Weldon BSc [99], Dr Esther Robinson PhD [64, 36], Prof Alan McNally PhD [41, 86], Dr Peter Muir PhD [64], Dr Ian B Vipond PhD [64], Dr John BoYes MBChB [29], Dr Venkat Sivaprakasam PhD [46], Dr Tranprit Saluja FRCPath/MD [75], Dr Samir Dervisevic FRCPath [54] and Dr Emma J Meader FRCPath [54].

**Leadership and supervision, and Sequencing and analysis:**
Dr Naomi R Park PhD [99], Karen Oliver BSc [99], Dr Aaron R Jeffries Ph.D. [91], Dr Sascha Ott PhD [94], Dr Ana da Silva Filipe PhD [48], Dr David A Simpson PhD [72] and Dr Chris Williams MB BS [69].

**Leadership and supervision, and Visualisation:**
Dr Jane AH Masoli MBChB [73, 91].

**Metadata curation, and Samples and logistics:**
Dr Bridget A Knight PhD. [73, 91], Dr Christopher R Jones Ph.D. [73, 91], Mr Cherian Koshy MSc CSci FIBMS [1], Miss Amy Ash BSc [1], Dr Anna Casey PhD [71], Dr Andrew Bosworth PhD [64, 36], Dr Liz Ratcliffe PhD [71], Dr Li Xu-McCrae PhD [36], Miss Hannah M Pymont MSc [64], Ms Stephanie Hutchings [64], Dr Lisa Berry PhD [84], Ms Katie Jones MSc [84], Dr Fenella Halstead PhD [46], Mr Thomas Davis MSc [21], Dr Christopher Holmes PhD [16], Prof Miren Iturriza-Gomara

PhD [92], Dr Anita O Lucaci PhD [92], Dr Paul Anthony Randell MBBCh [38, 104], Dr Alison Cox PhD [38, 104], Pinglawathee Madona [38, 104], Dr Kathryn Ann Harris PhD [30], Dr Julianne Rose Brown PhD [30], Dr Tabitha W Mahungu FRCPath [74], Dr Dianne Irish-Tavares FRCPath [74], Dr Tanzina Haque FRCPath PhD [74], Dr Jennifer Hart MRCP [74], Mr Eric Witele MSc [74], Mrs Melisa Louise Fenton DipHE [75], Mr Steven Liggett [79], Dr Clive Graham MD [56], Ms Emma Swindells Bsc [57], Ms Jennifer Collins BSc [50], Mr Gary Eltringham BSc [50], Ms Sharon Campbell MSc [17], Dr Patrick C McClure PhD [97], Dr Gemma Clark PhD [15], Dr Tim J Sloan PhD [60], Mr Carl Jones [15] and Dr Jessica Lynch PhD MBChB [2, 111].

**Metadata curation, and Sequencing and analysis:**
Dr Ben Warne MRCP [8], Steven Leonard PhD [99], Jillian Durham BSc [99], Dr Thomas Williams MD [90], Dr Sam T Haldenby PhD [92], Dr Nathaniel Storey PhD [30], Dr Nabil-Fareed Alikhan PhD [70], Dr Nadine Holmes PhD [18], Dr Christopher Moore PhD [18], Mr Matthew Carlile BSc [18], Malorie Perry MSc [69], Dr Noel Craine DPhil [69], Prof Ronan A Lyons MD [80], Miss Angela H Beckett MSc [13], Salman Goudarzi PhD [77], Christopher Fearn MRes [77], Kate Cook [77], Hannah Dent BSc [77] and Hannah Paul MRes [77].

**Metadata curation, and Software and analysis tools:**
Robert Davies [99].

**Project administration, and Samples and logistics:**
Beth Blane BSc [88], Sophia T Girgis MSc [88], Dr Mathew A Beale PhD [99], Katherine L Bellis [99, 88], Matthew J Dorman [99], Eleanor Drury [99], Leanne Kane [99], Sally Kay [99], Dr Samantha McGuigan [99], Dr Rachel Nelson PhD [99], Liam Prestwood [99], Dr Shavanthi Rajatileka PhD [99], Dr Rahul Batra MD [12], Dr Rachel J Williams PhD [82], Dr Mark Kristiansen PhD [82], Dr Angie Green PhD [98], Miss Anita Justice MSc [59], Dr Adhyana I.K Mahanama MD [81, 102] and Dr Buddhini Samaraweera MD [81, 102].

**Project administration, and Sequencing and analysis:**
Dr Nazreen F Hadjirin PhD [88] and Dr Joshua Quick PhD [41].

**Project administration, and Software and analysis tools:**
Mr Radoslaw Poplawski BSc [41].

**Samples and logistics, and Sequencing and analysis:**
Leanne M Kermack MSc [88], Nicola Reynolds PhD [7], Grant Hall BS [24], Yasmin Chaudhry BSc [24], Malte L Pinckert MPhil [24], Dr Iliana Georgana PhD [24], Dr Robin J Moll PhD [99], Dr Alicia Thornton [66], Dr Richard Myers [66], Dr Joanne Stockton PhD [41], Miss Charlotte A Williams BSc [82], Dr Wen C Yew PhD [58], Alexander J Trotter MRes [70], Miss Amy Trebes MSc [98], Mr George MacIntyre-Cockett BSc [98], Alec Birchley MSc [69], Alexander Adams BSc [69], Amy Plimmer [69], Bree Gatica-Wilcox MPhil [69], Dr Caoimhe McKerr PhD [69], Ember Hilvers MA [69], Hannah Jones [69], Dr Hibo Asad PhD [69], Jason Coombes BSc [69], Johnathan M Evans MSc [69], Laia Fina [69], Lauren Gilbert A-Levels [69], Lee Graham BSc [69], Michelle Cronin [69], Sara Kumziene-SummerhaYes MSc [69], Sarah Taylor [69], Sophie Jones MSc [69], Miss Danielle C Groves BA [93], Mrs Peijun Zhang MSc [93], Miss Marta Gallis MSc [93] and Miss Stavroula F Louka MSc [93].

**Samples and logistics, and Software and analysis tools:**
Dr Igor Starinskij Msc MRCP [48].

**Sequencing and analysis, and Software and analysis tools:**
Dr Chris J Illingworth PhD [47], Dr Chris Jackson PhD [47], Ms Marina Gourtovaia MSc [99], Gerry Tonkin-Hill [99], Kevin Lewis [99], Dr Jaime M Tovar-Corona PhD [99], Dr Keith James PhD [99], Dr Laura Baxter PhD [94], Dr Mohammad T. Alam PhD [94], Dr Richard J Orton PhD [48], Dr Joseph Hughes PhD [48], Dr Sreenu Vattipally PhD [48], Dr Manon Ragonnet-Cronin PhD [39], Dr Fabricia F. Nascimento PhD [39], Mr David Jorgensen MSc [39], Ms Olivia Boyd MSc [39], Ms Lily Geidelberg MSc [39], Dr Alex E Zarebski PhD [23], Dr Jayna Raghwani PhD [23], Dr Moritz UG Kraemer DPhil [23], Joel Southgate MSc [10, 69], Dr Benjamin B Lindsey MRCP [93] and Mr Timothy M Freeman MPhil [93].

**Software and analysis tools, and Visualisation:**
Jon-Paul Keatley [99], Dr Joshua B Singer PhD [48], Leonardo de Oliveira Martins PhD [70], Dr Corin A Yeats PhD [14], Dr Khalil Abudahab PhD [14, 114], Mr Ben EW Taylor MEng [14, 114] and Mirko Menegazzo [14].

**Leadership and supervision:**
Prof John Danesh [99], Wendy Hogsden MSc [46], Dr Sahar Eldirdiri MBBS MSc FRCPath [21], Mrs Anita Kenyon MSc [21], Dr Jenifer Mason MBBS [43], Mr Trevor I Robinson MSc [43], Prof Alison Holmes MD [38, 103], Dr James Price PhD [38, 103], Prof John A Hartley PhD [82], Dr Tanya Curran PhD [3], Dr Alison E Mather PhD [70], Dr Giri Shankar [69], Dr Rachel Jones [69], Dr Robin Howe [69] and Dr Sian Morgan FRCPath [9].

**Metadata curation:**
Dr Elizabeth Wastenge MD [53], Dr Michael R Chapman PhD [34, 88, 99], Mr Siddharth Mookerjee MPH [38, 103], Dr Rachael Stanley PhD [54], Mrs Wendy Smith [15], Prof Timothy Peto PhD [59], Dr David Eyre PhD [59], Dr Derrick Crook [59], Dr Gabrielle Vernet MBBS [33], Dr Christine Kitchen PhD [10], Huw Gulliver [10], Dr Ian Merrick PhD [10], Prof Martyn Guest PhD [10], Robert Munn BSc [10], Dr Declan T Bradley [63, 72] and Dr Tim Wyatt [63].

**Project administration:**
Dr Charlotte Beaver [99], Luke Foulser [99], Sophie Palmer [88], Carol M Churcher [88], Ellena Brooks MA [88], Kim S Smith [88], Dr Katerina Galai PhD [88], Georgina M McManus BSc [88], Dr Frances Bolt PhD [38, 103], Dr Francesc Coll PhD [19], Lizzie Meadows MA [70], Dr Stephen W Attwood PhD [23], Dr Alisha Davies [69], Elen De Lacy MSc [69], Fatima Downing [69], Sue Edwards [69], Dr Garry P Scarlett PhD [76], Mrs Sarah Jeremiah MSc [83] and Dr Nikki Smith PhD [93].

**Samples and logistics:**
Danielle Leek Bsc [88], Sushmita Sridhar BS [88, 99], Sally Forrest BSc [88], Claire Cormie [88], Harmeet K Gill PhD [88], Joana Dias MSc [88], Ellen E Higginson PhD [88], Mailis Maes MPhil [88], Jamie Young BSc [88], Michelle Wantoch PhD [7], Sanger Covid Team (www.sanger.ac.uk/covid-team) [99], Dorota Jamrozy [99], Stephanie Lo [99], Dr Minal Patel PhD [99], Verity Hill [90], Ms Claire M Bewshea MSc [91], Prof Sian Ellard FRCPath [73, 91], Dr Cressida Auckland FRCPath [73], Dr Ian Harrison [66], Dr Chloe Bishop [66], Dr Vicki Chalker [66], Dr Alex Richter PhD [85], Dr Andrew Beggs PhD [85], Dr Angus Best PhD [86], Dr Benita Percival PhD [86], Dr Jeremy Mirza PhD [86], Dr Oliver Megram PhD [86], Dr Megan Mayhew PhD [86], Dr Liam Crawford PhD [86], Dr Fiona Ashcroft PhD [86], Dr Emma Moles-Garcia PhD [86], Dr Nicola Cumley PhD [86], Mr Richard Hopes [64], Dr Patawee Asamaphan PhD [48], Mr Marc O Niebel MSc [48], Prof Rory N Gunson PhD FRCPath [100], Dr Amanda Bradley PhD [52], Dr Alasdair Maclean PhD [52], Dr Guy Mollett MBChB [52], Dr Rachel Blacow MBChB [52], Mr Paul Bird MSc [16], Mr Thomas Helmer [16], Miss Karlie Fallon [16], Dr Julian Tang [16], Dr Antony D Hale MBBS [49], Dr Louissa R Macfarlane-Smith PhD [49], Katherine L Harper MBiol [49], Miss Holli Carden MSc [49], Dr Nicholas W Machin MSc [45, 64], Ms Kathryn A Jackson MSc [92], Dr Shazaad S Y Ahmad MSc [45, 64], Dr Ryan P George PhD [45], Dr Lance Turtle PhD MRCP [92], Mrs Elaine O'Toole BSc [43], Mrs Joanne Watts BSc [43], Mrs Cassie Breen BSc [43], Mrs Angela Cowell MSc [43], Ms Adela Alcolea-Medina [32, 96], Ms Themoula Charalampous MSc [12, 42], Amita Patel [11], Dr Lisa J Levett PhD [35], Dr Judith Heaney PhD [35], Dr Aileen Rowan PhD [39], Prof Graham P Taylor DSc [39], Dr Divya Shah PhD [30], Miss Laura Atkinson MSc [30], Mr Jack CD Lee MSc [30], Mr Adam P Westhorpe BSc [82], Dr Riaz Jannoo PhD [82], Dr Helen L Lowe PhD [82], Miss Angeliki Karamani MSc [82], Miss Leah Ensell BSc [82], Mrs Wendy Chatterton MSc [35], Miss Monika Pusok MSc [35], Mrs Ashok Dadrah MSc [75], Miss Amanda Symmonds MSc [75], Dr Graciela Sluga MD/MSC [44], Dr Zoltan Molnar PhD [72], Mr Paul Baker MD [79], Prof Stephen Bonner [79], Ms Sarah Essex [79], Dr Edward Barton MD [56], Ms Debra Padgett BSc [56], Ms Garren Scott BSc [56], Ms Jane Greenaway MSc [57], Dr Brendan AI Payne MD [50], Dr Shirelle Burton-Fanning MD [50], Dr Sheila Waugh MD [50], Dr Veena Raviprakash MD [17], Ms Nicola Sheriff BSc [17], Ms Victoria Blakey BSc [17], ms Lesley-Anne Williams BSc [17], Dr Jonathan Moore MD [27], Ms Susanne Stonehouse BSc [27], Dr Louise Smith [55], Dr Rose K Davidson PhD [89], Dr Luke Bedford [26], Dr Lindsay Coupland PhD [54], Ms Victoria Wright BSc [18], Dr Joseph G Chappell PhD [97], Dr Theocharis Tsoleridis PhD [97], Prof Jonathan Ball PhD [97], Mrs Manjinder Khakh [15], Dr Vicki M

Fleming PhD [15], Dr Michelle M Lister PhD [15], Dr Hannah C Howson-Wells PhD [15], Dr Louise Berry [15], Dr Tim Boswell [15], Dr Amelia Joseph [15], Dr Iona Willingham [15], Dr Nichola Duckworth [60], Dr Sarah Walsh [60], Dr Emma Wise PhD [2, 111], Dr Nathan Moore PhD [2, 111], Miss Matilde Mori BSc [2, 108, 111], Dr Nick Cortes MRCP FRCPath [2, 111], Dr Stephen Kidd PhD [2, 111], Dr Rebecca Williams BMBS [33], Laura Gifford MSc [69], Miss Kelly Bicknell [61], Dr Sarah Wyllie [61], Miss Allyson Lloyd [61], Mr Robert Impey MSc [61], Ms Cassandra S Malone MSc [6], Mr Benjamin J Cogger BSc [6], Nick Levene MSc [62], Lynn Monaghan [62], Dr Alexander J Keeley MRCP [93], Dr David G Partridge FRCP FRCPath [78, 93], Dr Mohammad Raza [78, 93], Dr Cariad Evans [78, 93] and Dr Kate Johnson [78, 93].

**Sequencing and analysis:**
Emma Betteridge BSc [99], Ben W Farr BSc [99], Scott Goodwin MSc [99], Dr Michael A Quail PhD [99], Carol Scott [99], Lesley Shirley MSc [99], Scott AJ Thurston BSc [99], Diana Rajan MSc [99], Dr Iraad F Bronner PhD [99], Louise Aigrain PhD [99], Dr Nicholas M Redshaw PhD [99], Dr Stefanie V Lensing PhD [99], Shane McCarthy [99], Alex Makunin [99], Dr Carlos E Balcazar PhD [90], Dr Michael D Gallagher PhD [90], Dr Kathleen A Williamson PhD [90], Thomas D Stanton BSc [90], Ms Michelle L Michelsen BSc [91], Ms Joanna Warwick-Dugdale BSc [91], Dr Robin Manley Ph.D. [91], Ms Audrey Farbos MSc [91], Dr James W Harrison Ph.D. [91], Dr Christine M Sambles Ph.D. [91], Dr David J Studholme PhD. [91], Dr Angie Lackenby [66], Dr Tamyo Mbisa [66], Dr Steven Platt [66], Mr Shahjahan Miah [66], Dr David Bibby [66], Dr Carmen Manso [66], Dr Jonathan Hubb [66], Dr Gavin Dabrera [66], Dr Mary Ramsay [66], Dr Daniel Bradshaw [66], Dr Ulf Schaefer [66], Dr Natalie Groves [66], Dr Eileen Gallagher [66], Dr David Lee [66], Dr David Williams [66], Dr Nicholas Ellaby [66], Hassan Hartman [66], Nikos Manesis [66], Vineet Patel [66], Juan Ledesma [67], Ms Katherine A Twohig [67], Dr Elias Allara [64, 88], Ms Clare Pearson [64, 88], Mr Jeffrey K. J. Cheng MSc [94], Dr Hannah E. Bridgewater PhD [94], Ms Lucy R. Frost BSc [94], Ms Grace Taylor-Joyce BSc [94], Dr Paul E Brown PhD [94], Dr Lily Tong PhD [48], Ms Alice Broos BSc [48], Mr Daniel Mair BSc [48], Mrs Jenna Nichols BSc [48], Dr Stephen N Carmichael PhD [48], Dr Katherine L Smollett PhD [40], Dr Kyriaki Nomikou PhD [48], Dr Elihu Aranday-Cortes PhD/DVM [48], Ms Natasha Johnson BSc [48], Dr Seema Nickbakhsh PhD [48, 68], Dr Edith E Vamos PhD [92], Dr Margaret Hughes PhD [92], Dr Lucille Rainbow PhD [92], Mr Richard Eccles MSc [92], Ms Charlotte Nelson MSc [92], Dr Mark Whitehead PhD [92], Dr Richard Gregory PhD [92], Mr Matthew Gemmell MSc [92], Ms Claudia Wierzbicki BSc [92], Ms Hermione J Webster BSc [92], Ms Chloe L Fisher MSc [28], Mr Adrian W Signell BSc [20], Dr Gilberto Betancor PhD [20], Mr Harry D Wilson BSc [20], Dr Gaia Nebbia PhD FRCPath [12], Dr Flavia Flaviani PhD [31], Mr Alberto C Cerda MSc [96], Ms Tammy V Merrill MSc [96], Rebekah E Wilson MSc [96], Mr Marius Cotic MSc [82], Miss Nadua Bayzid BSc [82], Dr Thomas Thompson PhD [72], Dr Erwan Acheson PhD [72], Prof Steven Rushton PhD [51], Prof Sarah O'Brien PhD [51], David J Baker BEng [70], Steven Rudder [70], Alp Aydin MSci [70], Dr Fei Sang PhD [18], Dr Johnny Debebe PhD [18], Dr Sarah Francois PhD [23], Dr Tetyana I Vasylyeva DPhil [23], Dr Marina Escalera Zamudio PhD [23], Mr Bernardo Gutierrez MSc [23], Dr Angela Marchbank BSc [10], Joshua Maksimovic FD [9], Karla Spellman FD [9], Kathryn McCluggage Msc [9], Dr Mari Morgan PhD [69], Robert Beer BSc [9], Safiah Afifi BSc [9], Trudy Workman HNC [10], William Fuller BSc [10], Catherine Bresner Bsc [10], Dr Adrienn Angyal PhD [93], Dr Luke R Green PhD [93], Dr Paul J Parsons PhD [93], Miss Rachel M Tucker MSc [93], Dr Rebecca Brown PhD [93] and Mr Max Whiteley PhD [93]

**Software and analysis tools:**
James Bonfield BSc [99], Dr Christoph Puethe [99], Mr Andrew Whitwham BSc [99], Jennifier Liddle [99], Dr Will Rowe PhD [41], Dr Igor Siveroni PhD [39], Dr Thanh Le-Viet PhD [70] and Amy Gaskin MSc [69].

**Visualisation:**
Dr Rob Johnson PhD [39].

**1** Barking, Havering and Redbridge University Hospitals NHS Trust, **2** Basingstoke Hospital, **3** Belfast Health & Social Care Trust, **4** Betsi Cadwaladr University Health Board, **5** Big Data Institute, Nuffield Department of Medicine, University of Oxford, **6** Brighton and Sussex University Hospitals NHS Trust, **7** Cambridge Stem Cell Institute, University of Cambridge, **8** Cambridge University Hospitals NHS Foundation Trust, **9** Cardiff and Vale University Health Board, **10** Cardiff University, **11** Centre for Clinical Infection & Diagnostics Research, St. Thomas' Hospital and Kings College London, **12** Centre for Clinical Infection and Diagnostics Research, Department of Infectious Diseases, Guy's and St Thomas' NHS Foundation Trust, **13** Centre for Enzyme Innovation, University of Portsmouth (PORT), **14** Centre for Genomic Pathogen Surveillance, University of Oxford, **15** Clinical Microbiology Department, Queens Medical Centre, **16** Clinical Microbiology, University Hospitals of Leicester NHS Trust, **17** County Durham and Darlington NHS Foundation Trust, **18** Deep Seq, School of Life Sciences, Queens Medical Centre, University of Nottingham, **19** Department of Infection Biology, Faculty of Infectious & Tropical Diseases, London School of Hygiene & Tropical Medicine, **20** Department of Infectious Diseases, King's College London, **21** Department of Microbiology, Kettering General Hospital, **22** Departments of Infectious Diseases and Microbiology, Cambridge University Hospitals NHS Foundation Trust; Cambridge, UK, **23** Department of Zoology, University of Oxford, **24** Division of Virology, Department of Pathology, University of Cambridge, **25** East Kent Hospitals University NHS Foundation Trust, **26** East Suffolk and North Essex NHS Foundation Trust, **27** Gateshead Health NHS Foundation Trust, **28** Genomics Innovation Unit, Guy's and St. Thomas' NHS Foundation Trust, **29** Gloucestershire Hospitals NHS Foundation Trust, **30** Great Ormond Street Hospital for Children NHS Foundation Trust, **31** Guy's and St. Thomas' BRC, **32** Guy's and St. Thomas' Hospitals, **33** Hampshire Hospitals NHS Foundation Trust, **34** Health Data Research UK Cambridge, **35** Health Services Laboratories, **36** Heartlands Hospital, Birmingham, **37** Hub for Biotechnology in the Built Environment, Northumbria University, **38** Imperial College Hospitals NHS Trust, **39** Imperial College London, **40** Institute of Biodiversity, Animal Health & Comparative Medicine, **41** Institute of Microbiology and Infection, University of Birmingham, **42** King's College London, **43** Liverpool Clinical Laboratories, **44** Maidstone and Tunbridge Wells NHS Trust, **45** Manchester University NHS Foundation Trust, **46** Microbiology Department, Wye Valley NHS Trust, Hereford, **47** MRC Biostatistics Unit, University of Cambridge, **48** MRC-University of Glasgow Centre for Virus Research, **49** National Infection Service, PHE and Leeds Teaching Hospitals Trust, **50** Newcastle Hospitals NHS Foundation Trust, **51** Newcastle University, **52** NHS Greater Glasgow and Clyde, **53** NHS Lothian, **54** Norfolk and Norwich University Hospital, **55** Norfolk County Council, **56** North Cumbria Integrated Care NHS Foundation Trust, **57** North Tees and Hartlepool NHS Foundation Trust, **58** Northumbria University, **59** Oxford University Hospitals NHS Foundation Trust, **60** PathLinks, Northern Lincolnshire & Goole NHS Foundation Trust, **61** Portsmouth Hospitals University NHS Trust, **62** Princess Alexandra Hospital Microbiology Dept., **63** Public Health Agency, **64** Public Health England, **65** Public Health England, Clinical Microbiology and Public Health Laboratory, Cambridge, UK, **66** Public Health England, Colindale, **67** Public Health England, Colindale, **68** Public Health Scotland, **69** Public Health Wales NHS Trust, **70** Quadram Institute Bioscience, **71** Queen Elizabeth Hospital, **72** Queen's University Belfast, **73** Royal Devon and Exeter NHS Foundation Trust, **74** Royal Free NHS Trust, **75** Sandwell and West Birmingham NHS Trust, **76** School of Biological Sciences, University of Portsmouth (PORT), **77** School of Pharmacy and Biomedical Sciences, University of Portsmouth (PORT), **78** Sheffield Teaching Hospitals, **79** South Tees Hospitals NHS Foundation Trust, **80** Swansea University, **81** University Hospitals Southampton NHS Foundation Trust, **82** University College London, **83** University Hospital Southampton NHS Foundation Trust, **84** University Hospitals Coventry and Warwickshire, **85** University of Birmingham, **86** University of Birmingham Turnkey Laboratory, **87** University of Brighton, **88** University of Cambridge, **89** University of East Anglia, **90** University of Edinburgh, **91** University of Exeter, **92** University of Liverpool, **93** University of Sheffield, **94** University of Warwick, **95** University of Cambridge, **96** Viapath, Guy's and St Thomas' NHS Foundation Trust, and King's College Hospital NHS Foundation Trust, **97** Virology, School of Life Sciences, Queens Medical Centre, University of Nottingham, **98** Wellcome Centre for Human Genetics, Nuffield Department of Medicine, University of Oxford, **99** Wellcome Sanger Institute, **100** West of Scotland Specialist Virology Centre, NHS Greater Glasgow and Clyde, **101** Department of Medicine, University of Cambridge, **102** Ministry of Health, Sri Lanka, **103** NIHR Health Protection Research Unit in HCAI and AMR, Imperial College London, **104** North West London Pathology, **105** NU-OMICS, Northumbria University, **106** University of Kent, **107** University of Oxford, **108** University of Southampton, **109** University of Southampton School of Health Sciences, **110** University of Southampton School of Medicine, **111** University of Surrey, **112** Warwick Medical School and Institute of Precision Diagnostics, Pathology, UHCW NHS Trust, **113** Wellcome Africa Health Research Institute Durban and **114** Wellcome Genome Campus.

### Supplementary Files

**Supplementary File S1**
Pillar 1 genome metadata

**Supplementary File S2**
Periscope canonical sgRNA counts for Pillar 1 samples

**Supplementary File S3**
Periscope canonical sgRNA counts for Pillar 2 samples

**Supplementary File S4**
Pillar 1 sample days since symptom onset

**Supplementary File S5**
Pillar 2 genome metadata

**Supplementary File S6**
Periscope noncanonical sgRNA counts for Pillar 1 samples. Note: some samples do not have any predicted noncanonical sgRNA and so will not appear in this file.

**Supplementary File S7**
R code to recreate figures and analyses

### Supplementary Tables

| PANGO Lineage | Public Health England nomenclature | NextStrain | Defining amino acid substitutions/deletions |
|---|---|---|---|
| B.1.177 | NA | 20A.EU1 | **S:** A222V |
| | | | **N:** A220V |
| B.1.1.7 | VOC-202012/01 | 20I/501Y.V1 | **ORF1ab:** T1001I, A1708D, I2230T, del3675–3677 SGF |
| | | | **S:** del69–70 HV, del144 Y, N501Y, A570D, D614G, P681H, T761I, S982A, D1118H |
| | | | **ORF8**: Q27stop, R52I, Y73C |
| | | | **N:** D3L, S235F |
| B.1.351 | VOC-202012/02 | 20H/501Y.V2 | **ORF1ab:** K1655N |
| | | | **E:** P71L |
| | | | **N:** T205I |
| | | | **S:** K417N, E484K, N501Y, D614G, A701V |
| P.1 | VOC-202101/02 | 20J/501Y.V3 | **Orf1ab**: S1188L, K1795Q |
| | | | **S**: L18F, T20N, P26S, D138Y, R190S, K417T, E484K, N501Y, H655Y, T1027I |
| | | | **Orf3a:** G174C |
| | | | **Orf8:** E92K |
| | | | **N**: P80R |
| | | | **Other:** del:11288:9 |
| P.2 | VUI-202101/01 | 20B/S.484K | **Orf1ab**: L3468V, L3930F |
| | | | **S**: E484K, V1176F |
| | | | **N**: A119S |

**Table S1 - Defining amino acid substitutions and deletions present in SARS-CoV-2 lineages B.1.177, B.1.1.7, B.1.351, P.1 and P.2**

| Testing Pillar | Total Samples > 90% genome coverage | Total Samples > 50k |
|---|---|---|
| **Pillar 1** | 4376 | 3856 |
| **Pillar 2** | 718 | 584 |
| **Total** | 5094 | 4440 |

**Table S2 - Dataset Summary**

| | S | E | M | N | ORF6 |
|---|---|---|---|---|---|
| **(Intercept)** | 69.20 * | 8.13 *** | 83.05 | 78.69 | 76.60 * |
| | (32.67) | (1.53) | (45.98) | (42.41) | (30.04) |
| **Lineage B.1.1** | -4.52 | -0.19 | 36.09 | 40.01 | 37.92 |
| | (37.88) | (1.77) | (53.31) | (49.10) | (34.83) |
| **Lineage B.1.1.7** | 298.02 *** | 9.54 *** | 586.67 *** | 416.73 *** | -7.11 |
| | (37.58) | (1.76) | (52.89) | (48.73) | (34.55) |
| **Lineage B.1.177** | 52.48 | 4.34 * | 585.87 *** | 149.91 ** | -8.46 |
| | (36.29) | (1.70) | (51.07) | (47.05) | (33.36) |
| **Days Since Symptom Onset** | 5.88 | 0.57 *** | 6.26 | 2.02 | 9.81 ** |
| | (3.60) | (0.17) | (5.07) | (4.65) | (3.31) |
| **N** | 935 | 935 | 935 | 934 | 935 |
| **R2** | 0.11 | 0.06 | 0.24 | 0.11 | 0.02 |

*** $p < 0.001$; ** $p < 0.01$; * $p < 0.05$.

**Table S3 - Linear regression model exploring impact of lineage (B.1, B.1.1, B.1.1.7 and B.1.177) and Days since symptom onset at sampling on sgRNA levels for S, E, M, N and ORF6.** No significant interactions between lineage and Days since symptom onset was observed. Reference lineage used B.1.

| | N* | ORF10 | ORF3a | ORF7a | ORF8 |
|---|---|---|---|---|---|
| **(Intercept)** | -0.09 | 0.01 | 1.95 | 3.15 | 0.84 |
| | (0.56) | (0.02) | (3.44) | (14.32) | (0.43) |
| **Lineage B.1.1** | 2.24 *** | 0.00 | 6.85 | 1.65 | -0.79 |
| | (0.65) | (0.02) | (3.99) | (16.60) | (0.50) |
| **Lineage B.1.1.7** | 3.94 *** | 0.03 | -0.12 | 16.58 | -0.33 |
| | (0.65) | (0.02) | (3.96) | (16.47) | (0.50) |
| **Lineage B.1.177** | 0.05 | 0.01 | 0.56 | -0.20 | -0.43 |
| | (0.62) | (0.02) | (3.82) | (15.91) | (0.48) |
| **Days Since Symptom Onset** | 0.03 | -0.00 | -0.29 | 0.39 | 0.03 |
| | (0.06) | (0.00) | (0.38) | (1.58) | (0.05) |
| **N** | 935 | 935 | 935 | 935 | 935 |
| **R2** | 0.07 | 0.00 | 0.01 | 0.00 | 0.00 |

*** $p < 0.001$; ** $p < 0.01$; * $p < 0.05$.

**Table S4 - Linear regression model exploring impact of lineage (B.1, B.1.1, B.1.1.7 and B.1.177) and Days since symptom onset at sampling on sgRNA levels for ORF3a, ORF7a, ORF8 and ORF10.** N* refers to novel non-canonical sgRNA due to N R203K/G204R substitutions and present in all lineages containing these mutations (*17*). No significant interactions between lineage and Days since symptom onset was observed. Reference lineage used B.1.

### Supplementary Figures

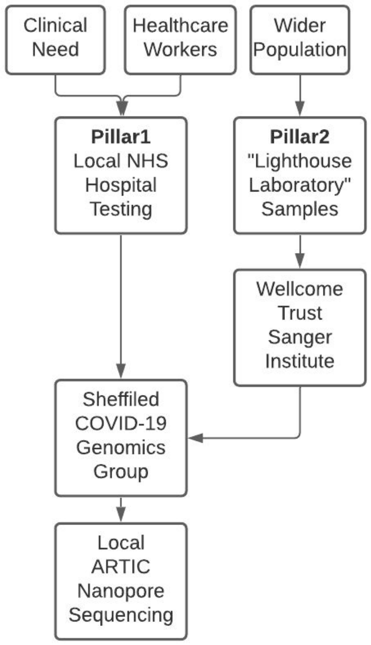


**Figure S1  - UK Diagnostic “Pillar” Testing Strategy**

Pillar 1: Diagnostic samples tested at Sheffield Teaching Hospitals NHS Foundation Trust, including patients and healthcare workers. Pillar 2: SARS-CoV-2 positive samples from the general public, tested at the UK’s “Lighthouse Laboratories” which are picked, and extracted at the Wellcome Trust Sanger Institute and sent to Sheffield COVID-19 Genomics Group for PCR, library preparation and sequencing.

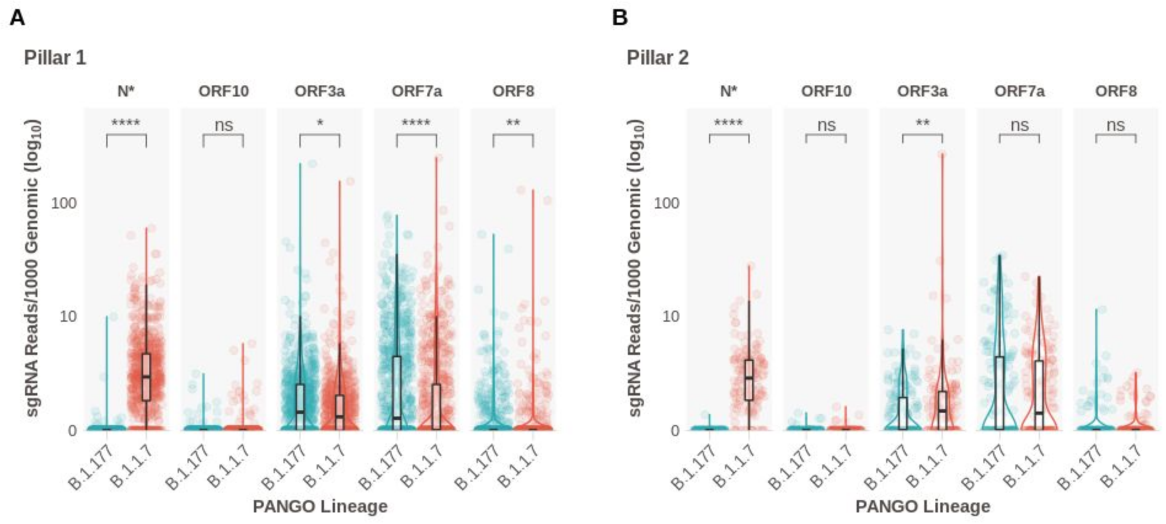


**Figure S2. Normalized sgRNA for ORF3a, ORF7a, ORF8 and ORF10**

**A.** Comparison of normalized sub-genomic RNA expression in B.1.177 and B.1.1.7 not plotted in main figures for Pillar 1 samples. **B.** Comparison of normalized sub-genomic RNA expression in B.1.177 and B.1.1.7 not plotted in main figures for Pillar 2 samples. N* refers to novel non-canonical sgRNA due to N R203K/G204R substitutions and present in all lineages containing these mutations (*17*).

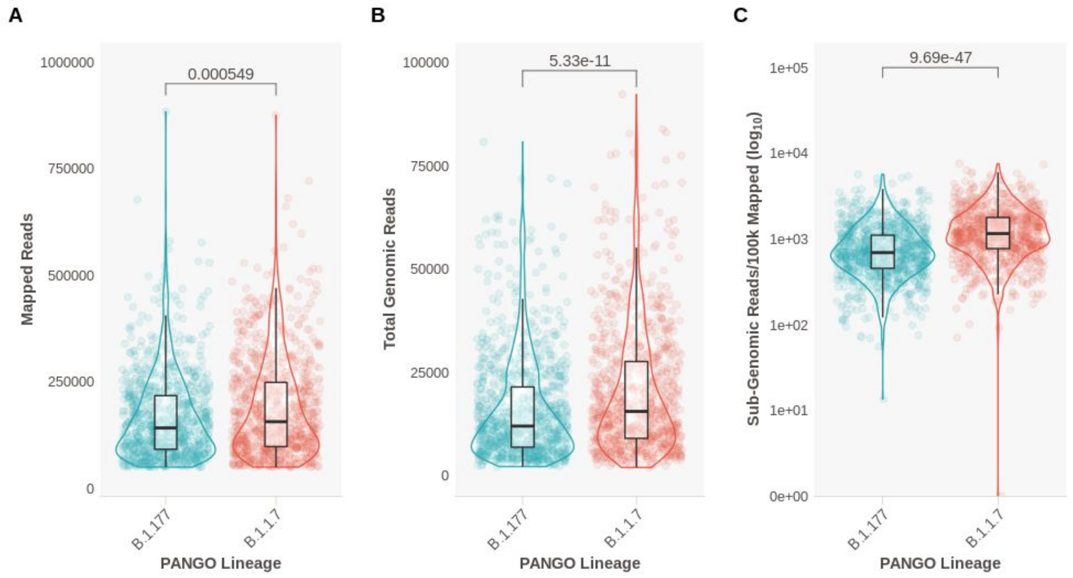


**Figure S3. Comparison of Mapped Reads and Genomic RNA (i.e. non-sgRNA) reads between B.1.177 and B.1.1.7 infections.**

**A.** Mapped reads between B.1.177 and B.1.1.7 (Wilcoxon effect size: 0.0810) **B.** Total genomic reads between B.1.177 and B.1.1.7 (Wilcoxon effect size: 0.154) **C.** Total sgRNA reads per 100,000 mapped reads (Wilcoxon effect size: 0.285) as an alternate normalization method to sgRNA normalization to genomic amplicon reads used in main figures.

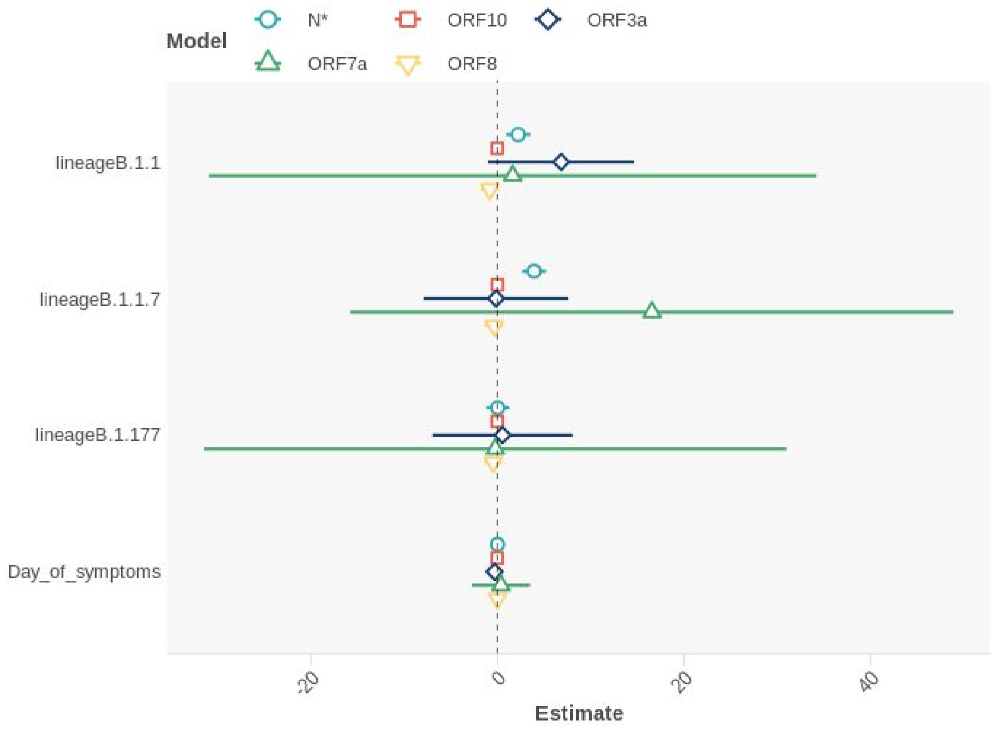


**Figure S4 - Linear regression model output evaluating impact of lineage and Days since symptom onset at sampling on ORF3a, ORF7a, ORF8 and ORF10.**

Effect size plot on the lower expressed ORFs in our dataset. Days since symptom onset has no effect. Reference lineage of B.1.

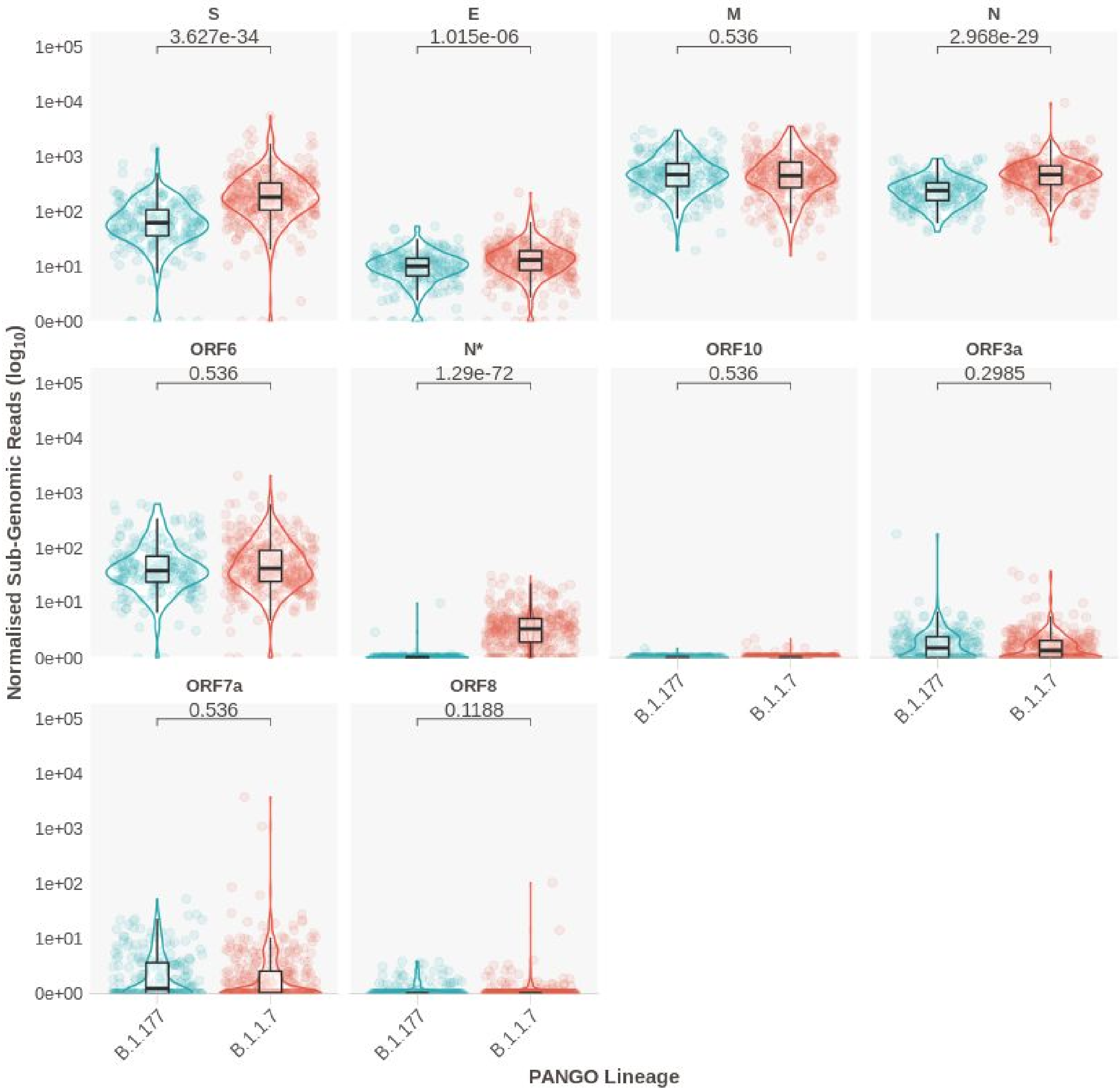


**Figure S5. Normalized sgRNA Expression in Healthcare Workers infected with B.1.177 and B.1.1.7 Lineages.**

Normalised sub-genomic expression stratified by lineage in only samples identified as originating from healthcare workers.

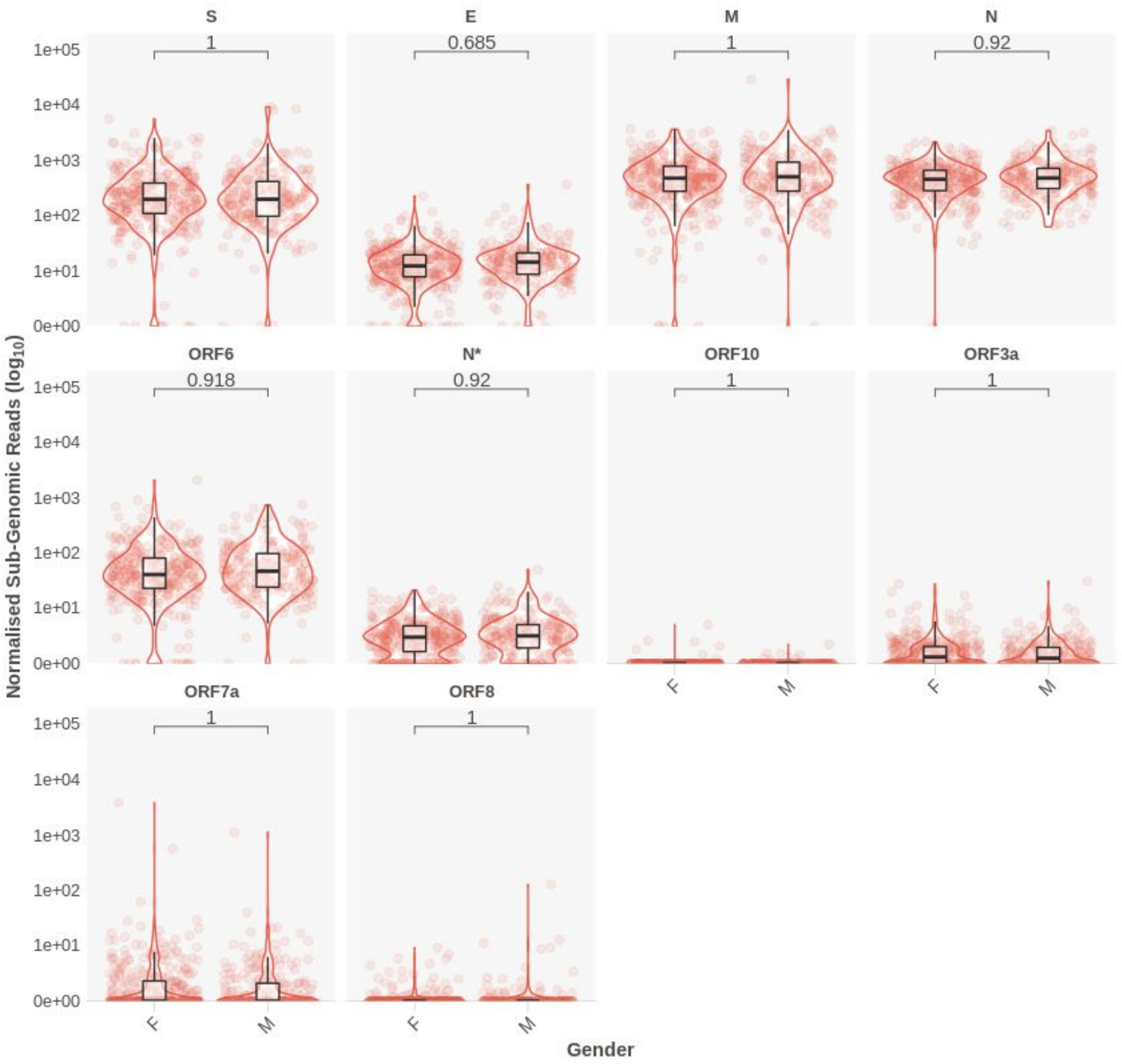


**Figure S6. Normalized sgRNA Expression in B.1.1.7 Stratified by Gender**

Normalized sub-genomic expression for samples of the B1.1.7 lineage by ORF and gender. No differences seen between infections in males and females. P values from a Wilcoxon unpaired test, adjusted for multiple testing with Holm. N* refers to novel non-canonical sgRNA due to N R203K/G204R substitutions and present in all lineages containing these mutations (*17*).

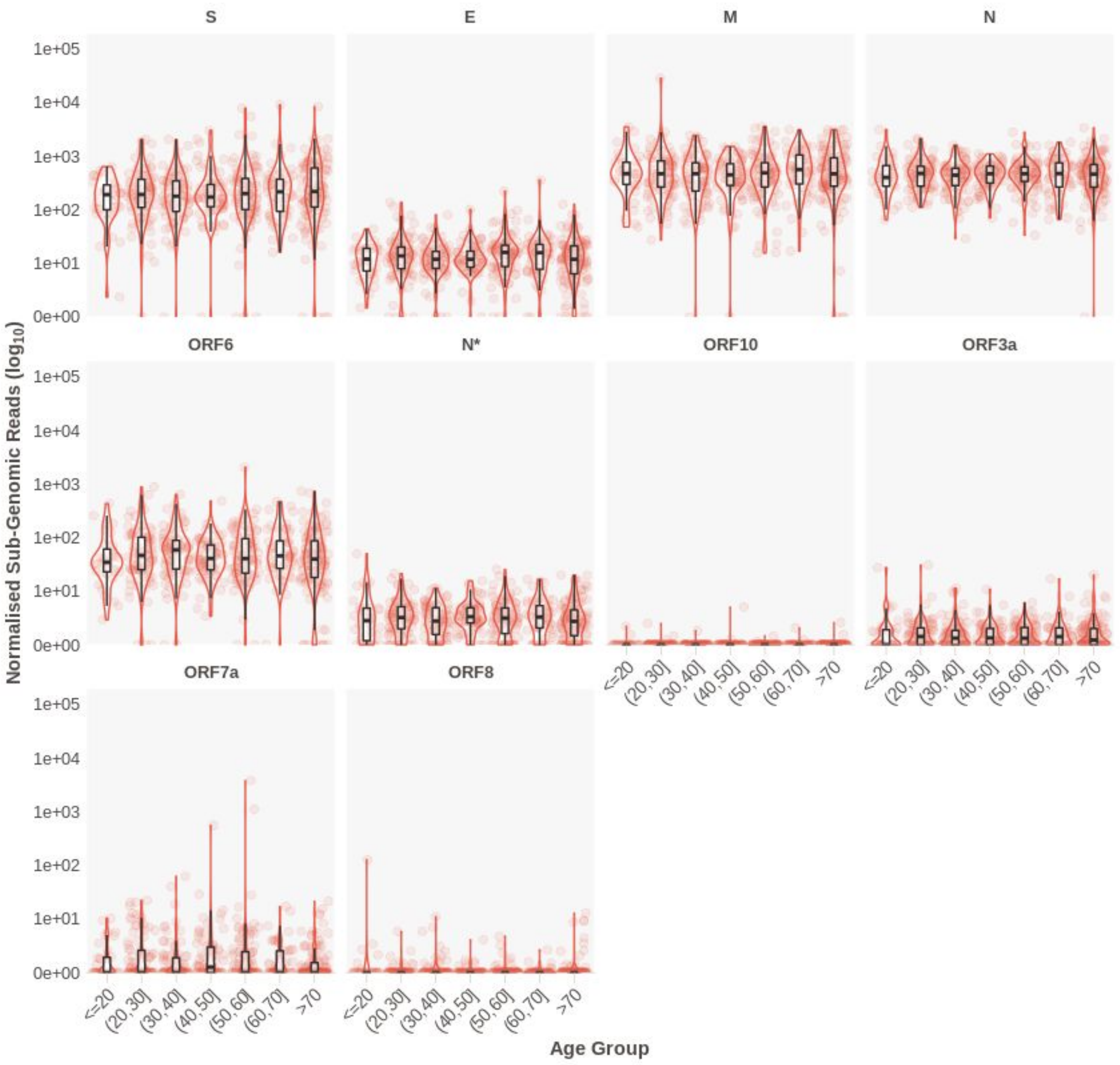


**Figure S7. Normalized sgRNA Expression in B.1.1.7 Stratified by Age**

Normalized sub-genomic expression for samples of the B.1.1.7 lineage by ORF and age group. X-axis notation is as follows; (a,b] means ">a and <=b". N* refers to novel non-canonical sgRNA due to N R203K/G204R substitutions and present in all lineages containing these mutations (*17*).

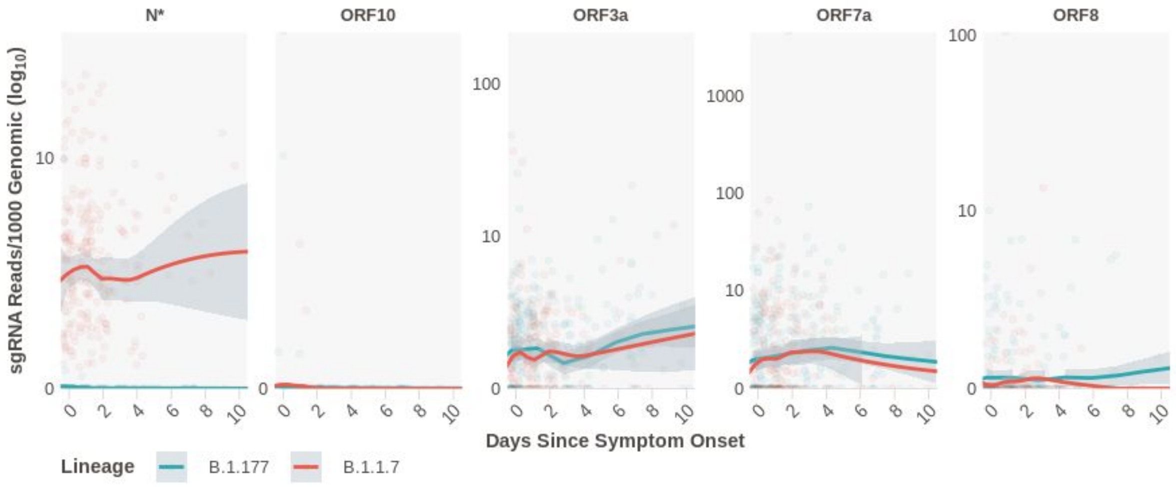


**Figure S8. Normalized sgRNA expression stratified by days since symptom onset for ORF3a, ORF7a, ORF8 and ORF10.** N* refers to novel non-canonical sgRNA due to N R203K/G204R substitutions and present in all lineages containing these mutations (*17*).

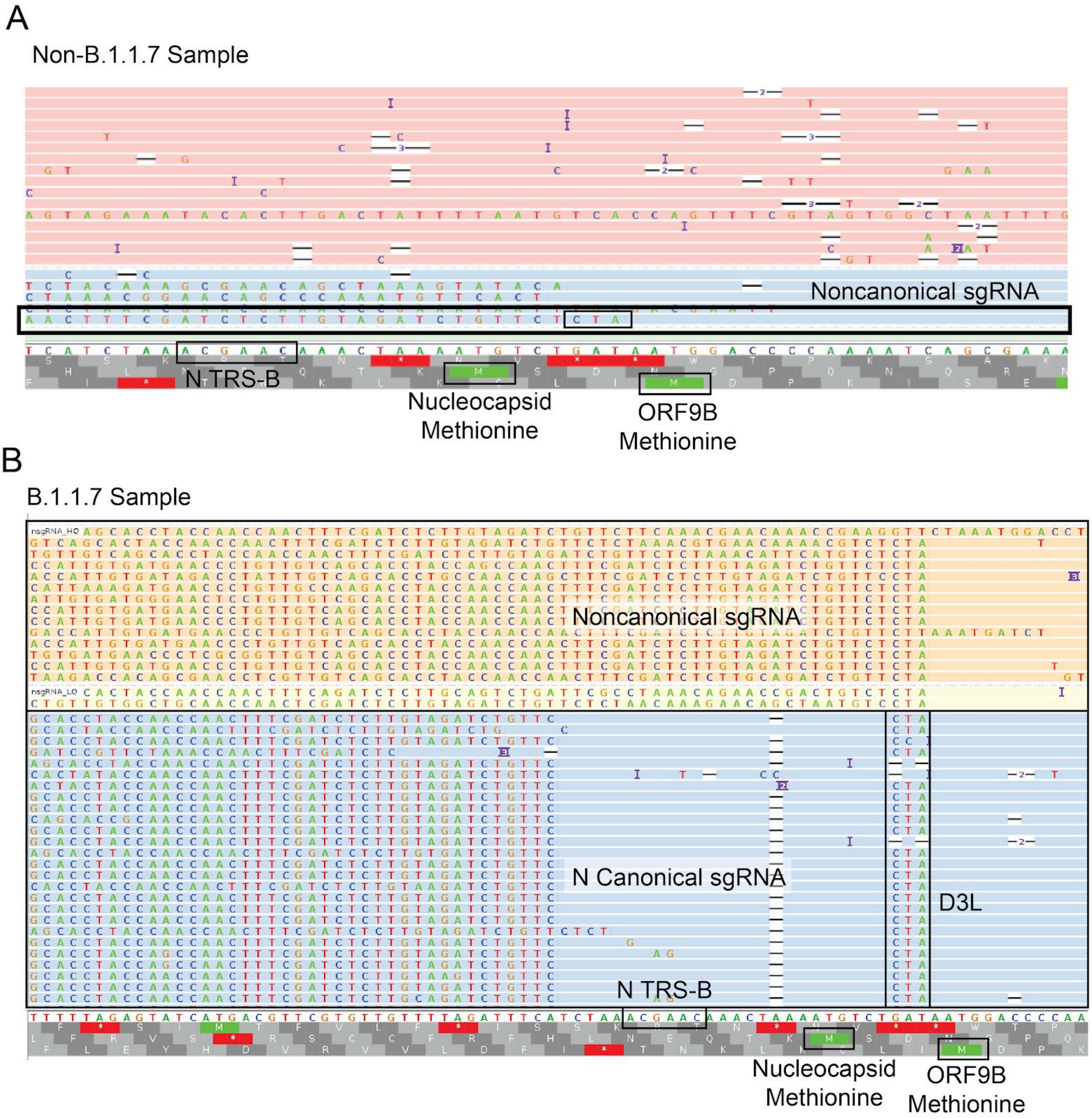


**Figure S9. Noncanonical sgRNA at nucleotide position 28282**

**A.** IGV plot showing reads around the canonical N TRS site in a non-B.1.1.7 sample. A Read is present that supports a 28282 noncanonical sgRNA that contains the CTA adjacent to the leader (red box). **B.** IGV plot showing reads around the canonical N TRS site in B.1.1.7 (navy bar bottom). Blue shaded reads are canonical ORF N sgRNA supporting reads, with the leader on the left. Peach reads are a noncanonical sgRNA produced from a new TRS like site complementary to a region 3’ of the leader sequence at the 5’ end of the genome as a result of the GAT->CTA mutation in this region.

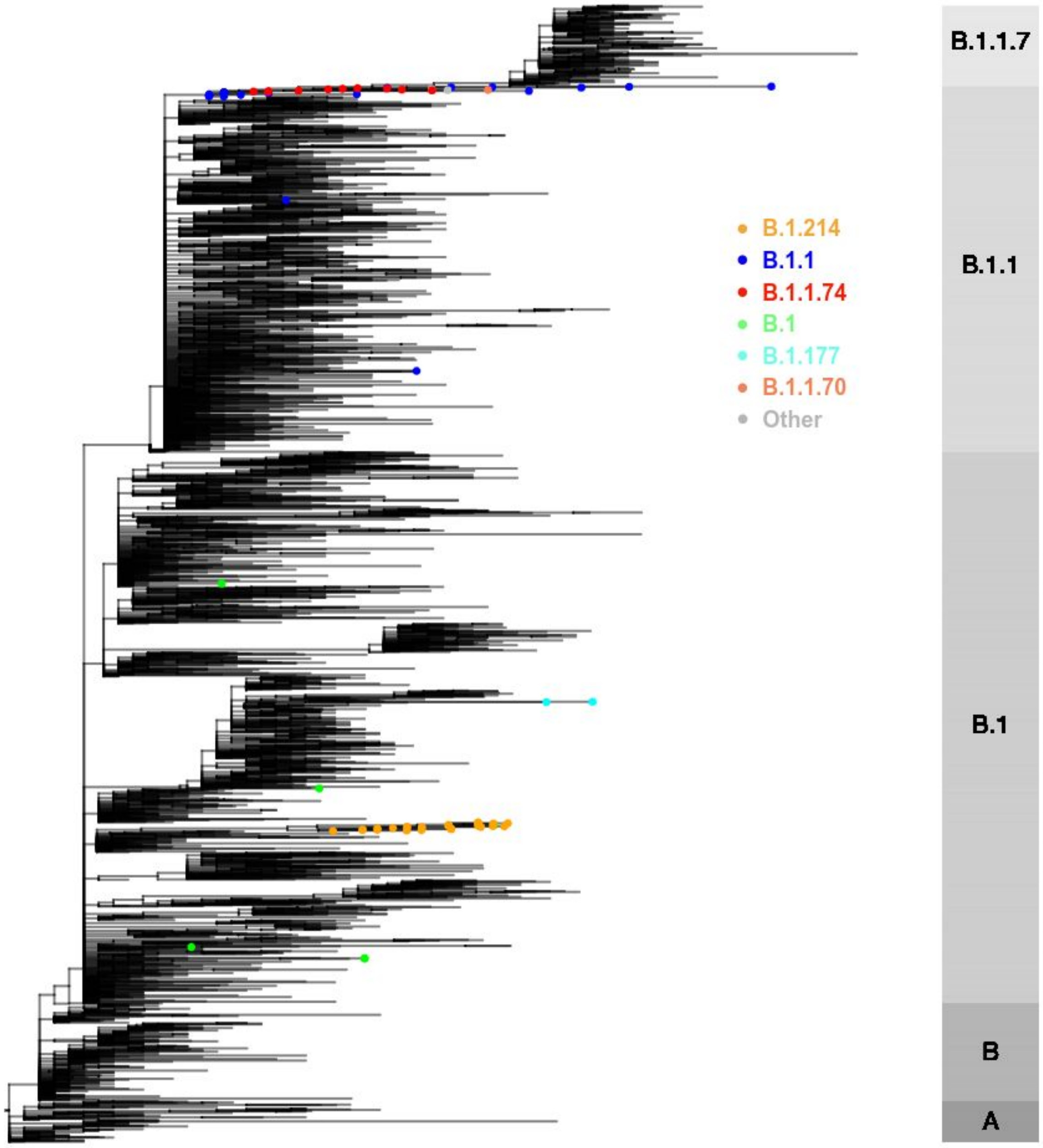


**Figure S10 - A representative phylogenetic tree of global SARS-CoV-2 genomes depicting non-B.1.1.7 viruses containing the nucleocapsid D3L mutation**

The nucleocapsid D3L mutation is found in 77 non-B.1.1.7 sequences indicated with dots at the ends of tips (see legend). Non-D3L sequences were subsampled for visualisation purposes. Bar on right depicts ancestral lineages alone. The grapevine pipeline (https://github.com/COG-UK/grapevine) was used for generating the phylogeny based on all data available on GISAID and COG-UK up until 16th February 2021. Of note, in some cases (e.g. B.1.1.74), several other B.1.1.7 defining mutations were also noted in sequences, raising the possibility of contamination, which could not be excluded.

A

| | ECT | RLU |
|---|---|---|
| (Intercept) | 25.1503877420282 *** | 1120.65777502961 *** |
| | (0.529559499520495) | (4.79528821880012) |
| relevel(lineage, ref = "B.1.177")B.1.1.7 | -0.86402311937367 | 44.9085071073261 *** |
| | (0.820756067729953) | (5.87090225491413) |
| Day_of_symptoms | 0.242207150335956 * | -0.973860685652852 |
| | (0.0982663834793713) | (1.03707256176755) |
| N | 142 | 579 |
| R2 | 0.0573069754451405 | 0.0990283849594999 |
| *** p < 0.001; ** p < 0.01; * p < 0.05. | | |

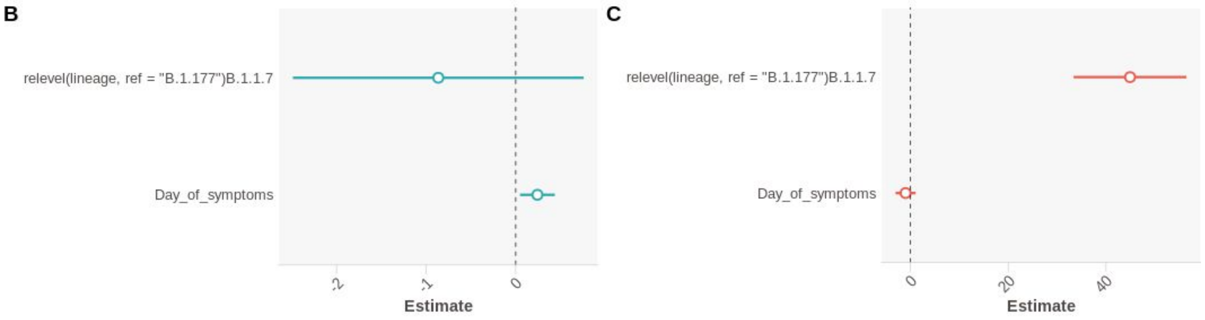


**Figure S11 - ECT and RLU are Most Influenced by Lineage**

**A.** Linear model of ECT and RLU by lineage and days since onset of symptoms. **B.** Effect plot of the linear model for ECT. **C.** Effect plot of the linear model for RLU. Reference lineage B.1.177 in both figures.
